# Marine community metabolomes carry fingerprints of phytoplankton community composition

**DOI:** 10.1101/2020.12.22.424086

**Authors:** Katherine R. Heal, Bryndan Durham, Angela K. Boysen, Laura T. Carlson, Wei Qin, François Ribalet, Angelicque E. White, Randelle M. Bundy, E. Virginia Armbrust, Anitra Eiding Ingalls

**Author notes:** Address correspondence to Anitra E. Ingalls,.

## Abstract

Phytoplankton transform inorganic carbon into thousands of biomolecules that represent an important pool of fixed carbon, nitrogen, and sulfur in the surface ocean. Metabolite production differs between phytoplankton, and the flux of these molecules through the microbial food web depends on compound-specific bioavailability to members of a wider microbial community. Yet relatively little is known about the diversity or concentration of metabolites within marine plankton. Here we compare 313 polar metabolites in 21 cultured phytoplankton species and in natural planktonic communities across environmental gradients to show that bulk community metabolomes reflect chemical composition of the phytoplankton community. We also show that groups of compounds have similar patterns across space and taxonomy suggesting that the concentrations of these compounds in the environment are controlled by similar sources and sinks. We quantify several compounds in the surface ocean that represent substantial understudied pools of labile carbon. For example, the N-containing metabolite homarine was up to 3% of particulate carbon and is produced in high concentrations by cultured *Synechococcus*, and S-containing gonyol accumulated up to 2.5 nM in surface particles and likely originates from dinoflagellates. Our results show that phytoplankton composition directly shapes the carbon composition of the surface ocean. Our findings suggest that in order to access these pools of bioavailable carbon, the wider microbial community must be adapted to phytoplankton community composition.

**IMPORTANCE:** Microscopic phytoplankton transform 100 million tons of inorganic carbon into thousands of different organic compounds each day. The structure of each chemical is critical to its biological and ecosystem function, yet, the diversity of biomolecules produced by marine microbial communities remained mainly unexplored, especially small polar molecules which are often considered the currency of the microbial loop. Here we explore the abundance and diversity of small biomolecules in planktonic communities across ecological gradients in the North Pacific and within 21 cultured phytoplankton species. Our work demonstrates that phytoplankton diversity is an important determinant of the chemical composition of the highly bioavailable pool of organic carbon in the ocean, and we highlight understudied yet abundant compounds in both the environment and cultured organisms. These findings add to understanding of how the chemical makeup of phytoplankton shapes marine microbial communities where the ability to sense and use biomolecules depends on the chemical structure.

## INTRODUCTION

In the ocean, the molecular makeup of organic carbon shapes its journey through the global carbon cycle. Phytoplankton fix approximately 100 million tons of carbon on a daily basis (1), roughly equivalent to half the total biomass of humans on earth (2). Each day, the microbial community respires about half of this carbon through the microbial loop (3). Approaches analyzing gene expression suggest freshly fixed small biomolecules, or metabolites, are among the most bioavailable in the surface ocean and represent a substantial conduit of carbon and energy flux. Much of the chemical complexity in phytoplankton-derived organic matter remains poorly described both qualitatively and quantitatively, particularly the highly labile portion of organic matter encompassing small polar metabolites. Here we characterize the small molecules within particulate organic matter in natural marine microbial communities in the North Pacific and cultures of 21 phytoplankton species to show that the chemical character of the bulk carbon pool in the ocean reflects the taxonomy of the primary producers present.

Small polar metabolites can be major carbon, nutrient, and/or energy sources for heterotrophs (4, 5) and are often considered the currency of the microbial loop in the ocean. Beyond this, they can maintain phytoplankton-bacterial interactions (6, 7), serve as micronutrients (8, 9, 10), manage redox stress (11), fuel nitrogen fixation (12), act as chemical defenses (13, 14), and more. The comprehensive analysis of the metabolites in a system (metabolomics) is a nascent field and analytically challenging in environmental settings (15, 16, 17). Metabolomic studies are being used to investigate physiological changes in marine organisms under laboratory conditions (4, 18, 19, 20, 21), though the same techniques have not been widely applied to whole communities in natural environments. Existing community marine metabolomic studies have employed targeted approaches in which the compounds detected are chosen by the analyst (5, 12, 21, 22, 23) or the analytical techniques employed preclude the observation of small, highly polar compounds (24, 25).

The chemical makeup of small polar compounds in freshly fixed organic matter influences the flux of carbon and energy through the microbial loop in the surface ocean. Here, we determine the metabolite pools in natural marine communities across space to explore the distributions of both known and unknown compounds. We compare our field observations to metabolomes of cultured marine primary producers from a broad taxonomic range and show how primary producers play an active role in shaping the chemical environment of the surface ocean. Finally, we highlight small polar compounds that may serve as potentially significant conduits of energy and nutrients in marine systems.

## RESULTS AND DISCUSSION

### Patterns of metabolites across space and taxonomy

We explored the patterns of metabolite abundances in three sample sets of marine particles in the North Pacific Ocean: one surface meridional transect and two depth profiles (Figure 1A, Table S1). In the transect sample set, seven general patterns emerged across latitude using a *k*-medoids clustering approach of 313 metabolites (Figure 1B). The most common pattern (40% of compounds) showed a modest yet robust increase in concentration with latitude (‘mode *a*’ in Figure 1B). This is likely related to the general increase in biomass with latitude (Figure S1 and as seen in the increase of chlorophyll in Figure 1A). Many compounds (30%) had their highest concentration in samples from 33 or 34°N (Figure 1B, ‘modes *c-f*’). About 19% of metabolites did not have a clear pattern with latitude (Figure 1B, ‘mode *b*’), while 8% of the compounds were generally more abundant in the southern samples than the northern samples (Figure 1 B, ‘mode *g*).

**FIG 1.**
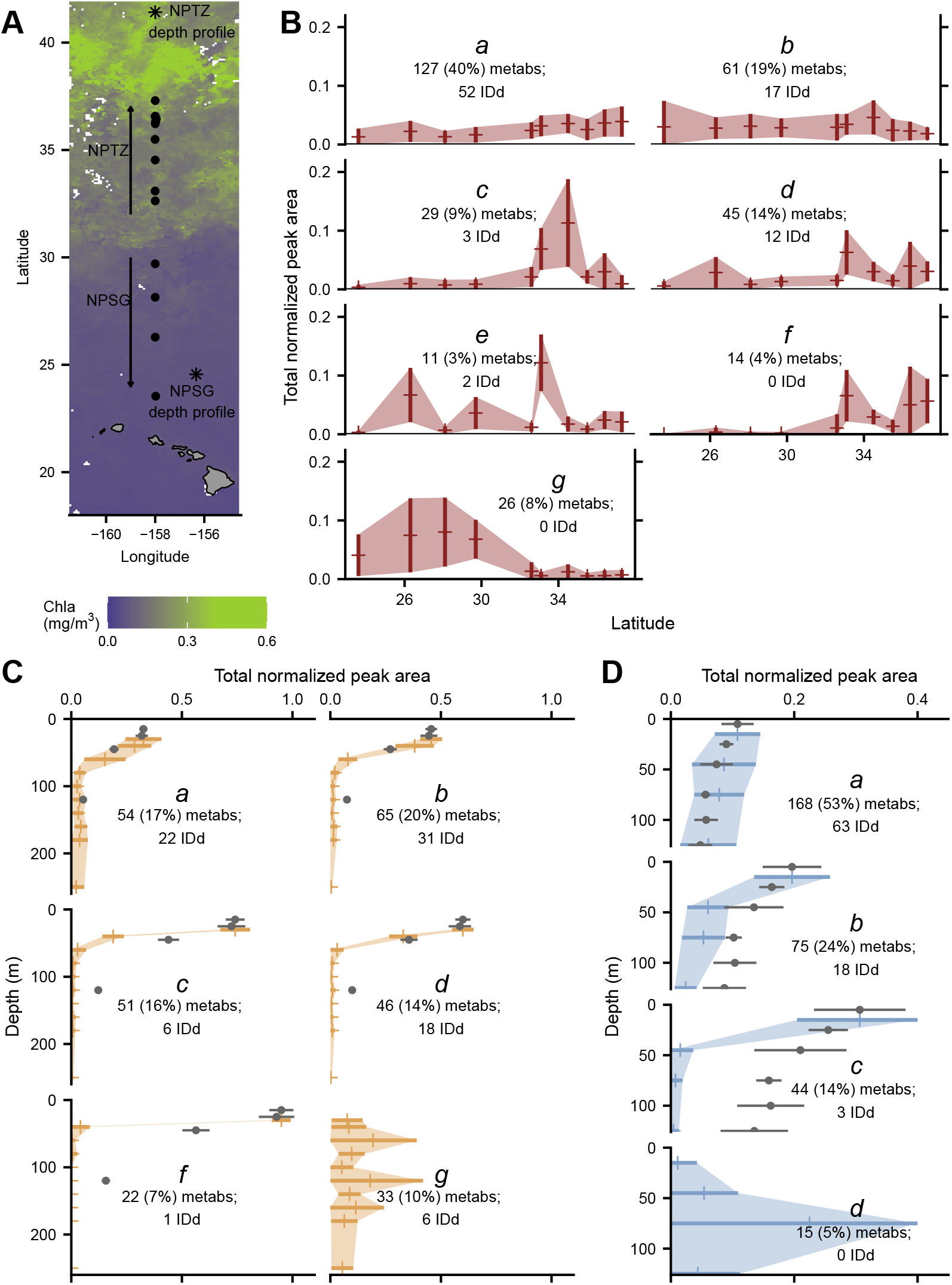
Sampling location and metabolite patterns. Map of sample locations of transect samples (dot) and depth profiles (asterisks). Map is overlayed with satellite-derived (MODIS-Aqua) chlorophyll at 8 day, 9 km resolution over the time period of the transect sampling (A). Patterns of total normalized metabolite concentrations found in each environmental dataset grouped into modes as a result of *k*-medoids clustering, plotted as one standard deviation around the mean (B, meridional transect; C, NPTZ depth profile; D, NPSG depth profile). We have excluded modes with fewer than 10 compounds in each dataset. Maximum normalized bulk PC is plotted over depth profiles, with surface PC concentration plotted to match surface total normalized metabolite peak area in order to compare the shape of attenuation (excluded in modes that do not attenuate with depth), grey dots with error bars (standard deviation, often smaller than markers). Number of metabolites (metabs) and percent of metabolites assigned to each cluster is noted, as well as the number of compounds identified (IDd) in each cluster. Full results are presented in Table S5 with cluster assignments in Table S6.

In a non-metric multidimensional scaling (NMDS) analysis where each metabolite was treated with equal weight, there was a distinct shift in metabolite patterns in the samples on either side of approximately 30°N (Figure S2, ANOSIM stat = 0.316, *p* = 0.005). This corresponds well with the southern boundary of the North Pacific Transition Zone, a well described oceanographic feature which extends from Japan to North America and arises from large-scale ocean circulation (26,27). The northern and southern edges of this transition zone comprise of rapid changes in thermohaline structure and biological species composition (28, 26, 29). We saw a similar stark transition within the metabolite pools that reiterate the transition from the warm, oligotrophic, North Pacific Subtropical Gyre (NPSG) into the colder, more nutrient replete North Pacific Transition Zone (NPTZ) where chlorophyll concentrations, *Synechococcus*, and picoeukaryote assemblages flourish (Figures 1 and S3, Table S2 for oceanographic conditions). Interestingly, the difference between metabolite profiles within the northern samples encompasses a much wider range in multivariate space, even within samples collected at the same time and location (biological replicates, Figure S2). This suggests the NPTZ is more heterogeneous in its metabolite profiles than the NPSG, and is supported by the observed high variability in particulate carbon (PC) at northern sampling sites (Figure S1).

We performed the same clustering technique (*k*-medoids) on the same metabolites (when observed) within two depth profiles: one from the NPSG and one from the NPTZ (Figure 1A for locations). Most of the metabolite concentrations decreased with depth, again corresponding with a decrease of PC (Figure 1C’modes a-f’and Figure 1D ‘modes *a-c*’). The extent of decrease in concentrations varied among metabolites, exemplified in the NPTZ depth profile by comparing the sharply attenuating ‘mode *c*’ to the more gentle attenuation of’mode *a*’(Figure 1C). Modes *a* in both NPTZ and NPSG depth profiles follow the PC pattern closely, but the other modes do not follow the same trend as bulk PC with depth (Figure 1C and D). A minority of compounds in both of these depth profiles either had a subsurface maximum or no clear relationship with depth (Figure 1C ‘mode *g* and Figure 1D ‘mode *d*’).

Using the 313 metabolites from the transect sample set as a template, we searched for the same compounds within metabolomes of 21 species of axenic phytoplankton grown under controlled conditions and analyzed on the same instrumental set up (5) (Tables S3 and S7). Phytoplankton are the primary source of fixed carbon to the surface ocean and our cultures were grown under conditions that support autotrophic growth so we could interrogate the metabolite pools these organisms produce *de novo* from inorganic components. The cultures explored here encompass a wide taxonomic range from picocyanobacteria that dominate much of our transect (Figure S3) to members of ubiquitous eukaryotic phytoplankton lineages like diatoms and coccolithophores. The taxonomic groups were recapitulated after a multivariate analysis of the metabolites across this data set in a semi-quantitative manner, using both NMDS (Figure S4) and *k*-medoids clustering (Figure 2A).

**FIG 2.**
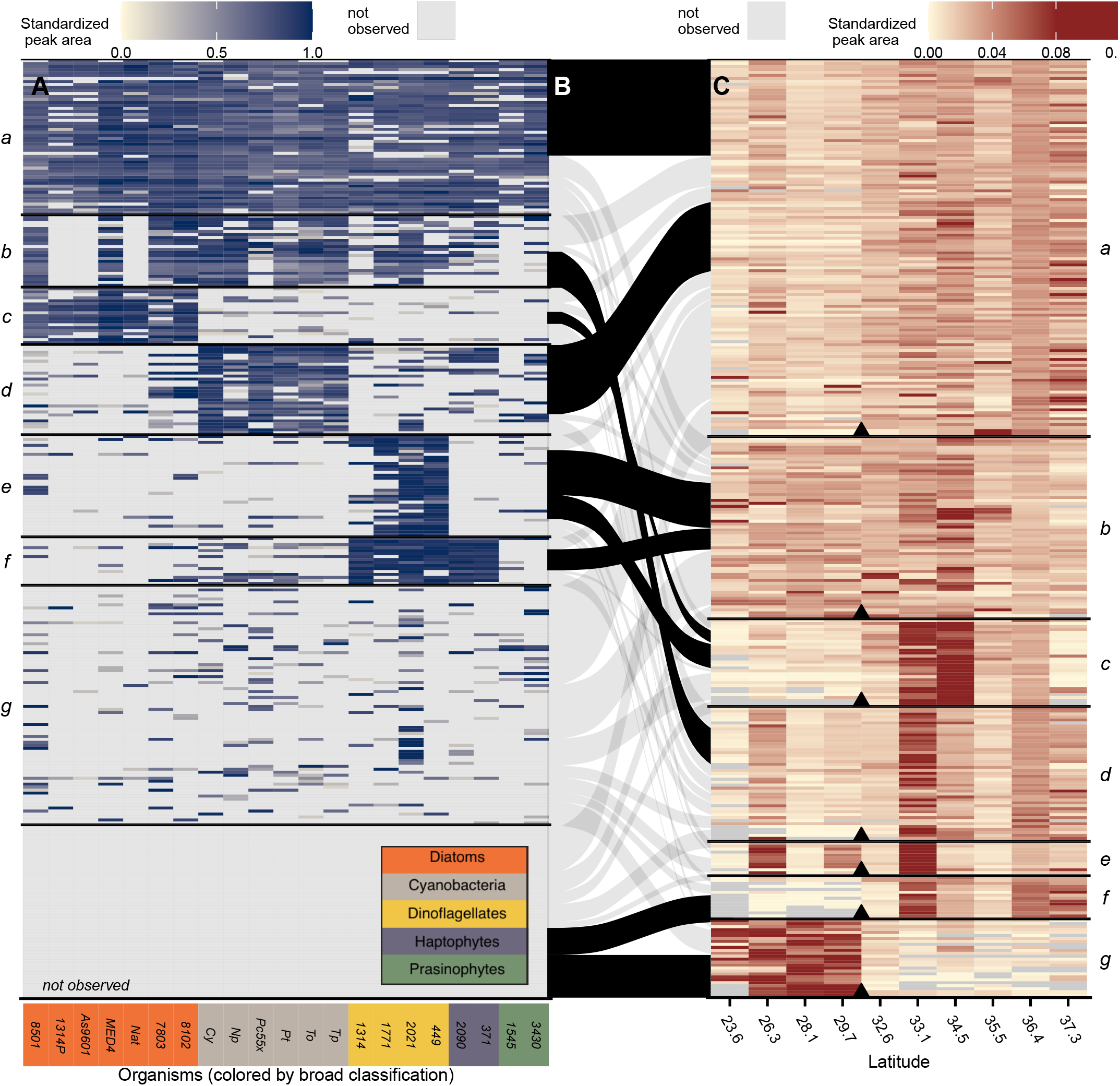
Relative abundance patterns in environmental and culture metabolites. Each row is a metabolite — either identified or unknown. Left (A) is the standardized peak areas for metabolites in the culture data sets. Right (C) is the relative abundance between samples along the meridional transect. Tile panels are grouped separately using a *k*-medoids clustering and reordered within each mode for visual clarity. The middle panel (B) shows which metabolites are shared between the culture and environmental *k*-medoids derived modes, with over-enriched connections between modes shown in black (*p* < 0.05 by bootstrap test) and remaining non-statistically significant connections shown in grey. Organisms are colored by broad taxonomic classification as shown in inset (orange = cyanobacteria, grey = diatoms, yellow = dinoflagellates, purple = haptophytes, green = prasinophytes).

Overall, we saw that 17% (52) of the 313 metabolites were present in most of the cultured organisms with 44 of the metabolites within this mode observed in over 80% of the phytoplankton species (‘mode *a*’ in Figure 2A). This suggests a set of compounds observable in most phytoplankton when analyzed under our analytical conditions. We were able to identify most of these compounds (33/52, 64%) which include many amino acids, primary metabolites, and nucleic acids (Tables S5 and S6). The remaining 36% of compounds within this mode could not be identified, demonstrating that even the compounds critical to the physiology and biochemistry of a broad swath of marine primary producers in natural systems remain elusive. Another 39% (123) of metabolites were seen primarily in subsets of organisms, separating into five modes (‘modes *b-f* in Figure 2A). Finally, about 44% of the 313 metabolites were either rarely or never observed in our cultures (mode *g* and *not observed* in Figure 2A).

The patterns of metabolites across the cultures suggest suites of compounds that are closely associated with taxonomic groups of organisms. Several identified metabolites in these groups corroborate previous work showing that certain types of organisms produce high concentrations of particular small molecules. For instance, DHPS and isethionate are within the mode of metabolites associated with diatoms (‘mode *d*’) (5, 7), taurine is associated with dinoflagellates and haptophytes (‘mode *f*’) (5), and glucosylglycerol is associated with cyanobacteria (‘mode *d*’) (30) (Figure 2A, Table S6). Most of the taxon-associated metabolites (72% of metabolites in modes *b-f*’ in Figure 2A) are still unidentified and offer possible future taxon-specific biomarkers in the polar organic carbon pools.

### Primary producers leave a metabolite signature in the environment

Compounds with similar patterns across these data sets would suggest shared sources and sinks. To assess this, we tested whether each *k*-medoids derived mode (within each sample set) was enriched in metabolites from a given mode from a different sample set, beyond what would be expected with a random assignment (assessed by a Monte Carlo-based bootstrapping approach, *p*-value < 0.05). For example, of the 52 metabolites that were observed in most of our cultured phytoplankton (‘mode *a*’in Figure 2A), there was a robust enrichment of compounds from the meridional transect ‘mode *a*’ (in Figure 2C, general increase with latitude, *p* < 0.01, Figure 2B). This may reflect a general increase in phytoplankton biomass with latitude as supported by the increase in PC and chlorophyll (Figures S1 and 1A).

We capitalized on our results to search for enrichment between each mode in each sample set and visualized enriched connections among all sample sets in a network analysis where each connection is a statistically significant enrichment between two modes (*p* < 0.05, Figure 3). This analysis revealed a few *meta-clusters*, or groups of compounds that have similar patterns as each other across different spatial and taxonomic ranges (Figure 3). These meta-clusters suggest compounds have similar sources across taxonomy that persist across both latitudinal and depth gradients in the environment. Building on the observation of over enrichment of metabolites between transect data set ‘mode *a*’ and culture data set ‘mode *a*’, we also saw that these modes share metabolites well beyond random assignment with the modes of metabolites in both depth profiles that attenuated in close proportion to PC with depth (‘modes *a*’ and *b* NPTZ depth profile, ‘mode *a*’ in NPSG depth profile, *p* < 0.05, Figure 3). Identified compounds within this ‘core metabolome’ meta-cluster included the amino acid glutamic acid, the nucleoside adenosine, the amino acid precursor homoserine, and several other primary metabolites (Table S6).

**FIG 3.**
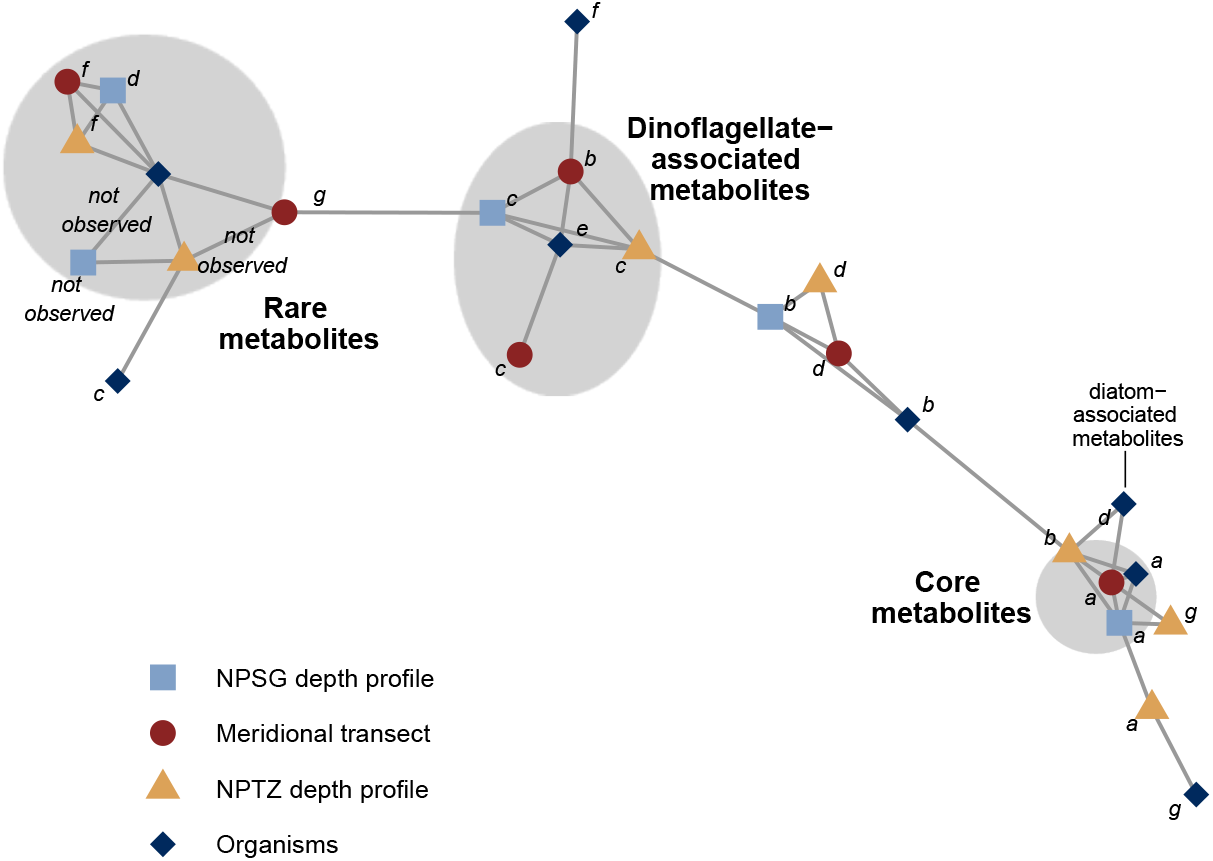
Network visualization of significant overlap between data sets shown in a network visualization. Each mode in each data set is depicted as a node (colored and shaped by data set, labeled as in Figures 1 and 2). Meta-clusters are highlighted in grey and labeled as described in text. Edges are connections between modes that are over-enriched in the same compounds (*p* < 0.05). Compound assignments to each mode and meta-cluster (if applicable) are found in Table S6.

Beyond the ‘core metabolome’, 77% of the 30 compounds associated tightly with diatoms (‘mode *d*’ in Figure 2A) were also over-represented within the group of compounds with a general increase with latitude (*p* < 0.01, Figure 2B). This pattern corresponded with an increase in fucoxanthin, a diatom biomarker observed to be increasing with latitude in a separate analysis from the same sampling period (Figure S6). The diatom-associated metabolites were also over-represented in the medium attenuating metabolites from the NPTZ depth profiles (‘mode *b*’ in Figure 1C and 3). DHPS (a probable osmolyte produced in high concentrations by diatoms (5, 21)) and glycerophosphocholine (a headgroup of phosphatidylcholine lipids commonly produced by eukaryotic phytoplankon in marine systems (31)) sat within this pattern space as did 11 other unidentified compounds (Table S6).

Surprisingly, compounds tightly associated with dinoflagellates (‘mode *e*’ in Figure 2A) showed a significant over-representation with an environmental distribution showing a distinct increase in concentration at 34.5°N (Figure 2). Metabolites displaying these patterns were over-represented in the sharply attenuating modes in the two depth profiles (dinoflagellate-associated meta-cluster in Figure 3), in contrast to the metabolites found associated with diatoms. None of the compounds that reside in this interaction space could be identified, leaving room for future work to identify and leverage these compounds as possible biomarkers for dinoflagellates that are easily observable in the environment.

Many of the compounds observed in our environmental samples but not observed in our culture dataset (‘not observed’ in Figure 2A) were over-represented by compounds that were more abundant in the NPSG than the NPTZ in the transect (‘mode *g* in Figure 2C) or increased with depth in the two depth profiles (rare metabolites metacluster in Figure 3). We only analyzed phytoplankton in our initial analysis; therefore it is likely that a subset of these compounds were produced *de novo* by organisms we did not survey. For instance, the compound *β*-glutamic acid was found to be more abundant at depth than in the surface waters in both of our depth profiles, in contrast to the majority of compounds observed (Figure S7) and was absent from our phytoplankton cultures (Table S5). *β*-glutamic acid is a major osmolyte in methanogenic archaea (32, 33), prompting us to search for this compound in *Nitrosopumilus maritimus* strain SCM1, a model species of Marine Group I Thaumarchaeota that are abundant in the ocean’s subsurface (34). We grew *N. maritimus*, analyzed its metabolome, and found *β*-glutamic acid as the most abundant identified metabolite, present at an intracellular concentrations of 730 mM (Table S6).

It is likely that some compounds in this group were not made ‘freshly by primary producers like phytoplankton or ammonia oxidizing archaea but were rather a signature of reworked particulate matter. For example, the compound arsenobetaine, which we detected in all of our environmental samples and, similar to *β*-glutamic acid, generally increased with depth in the depth profiles (Figure S7). This compound is a byproduct of heterotrophic degradation of phytoplankton-produced arseno-metabolites (35) and would therefore necessitate a co-culture in order to be observed in a laboratory setting (as well as a growth media with arsenic). Finally, it is likely that the cultures explored here were not producing all the compounds they are genetically able to produce — in previous laboratory experiments certain metabolites accumulate in cultures under specific environmental conditions and are not detectable under other environmental conditions (20, 21). If the production of certain compounds is variable or at rates below detection, we may not have seen them on our culture data.

### Metabolites as a quantitative component of the bulk carbon pool

We obtained absolute concentrations of the identified compounds to better understand the quantitative importance of these different metabolites within the particulate carbon landscape. The combined concentration of the identified metabolites (85 of the 313 total) ranged from 68-234 nM particulate carbon in the surface transect samples (Figures S1, and S8; Table S9). This corresponds to 2.9% (± 1.0%) to 5.2% (± 1.4%) of the particulate carbon pool and 2.6% (± 1.0%) to 8.2% (± 2.4%) of the particulate nitrogen pool across this transect (Table S9). There was no clear pattern in the percent of particulate carbon or nitrogen characterized by the quantifiable metabolites with latitude; this is likely confounded by the high variability in the particulate carbon and nitrogen measurements and the low geographical resolution of the metabolite sampling. In the NPTZ depth profile, we quantified 17-966 nM of particulate carbon in the metabolite pool, corresponding to a rough estimate of 10% of the particulate carbon and nitrogen pools in the surface sample (Table S9). In the NPSG profile, we quantified approximately 3.7% of the total carbon pool in the surface sample (Table S9). The concentration of surface particulate metabolites was approximately two times higher than what we observed a year later in the NPSG (during the transect sampling, Table S9), likely due to the fact that the NPSG depth profile was sampled within an anticyclonic eddy with high surface primary productivity and particulate carbon (36).

Quantitatively, the environmental metabolite pools were dominated by a handful of abundant compounds, similar to previous work (12, 23).There were obvious differences in metabolite composition between the three environmental samplings (Figure 4). For example, on a molar basis, glycine betaine contributed to up to 17% of the quantified metabolite pool in samples below 125 m in the NPTZ, substantially more (on a mole fraction basis) than the other data sets. In contrast, the NPSG depth profile had high contributions from gonyol in the surface and guanine at depth.

**FIG 4.**
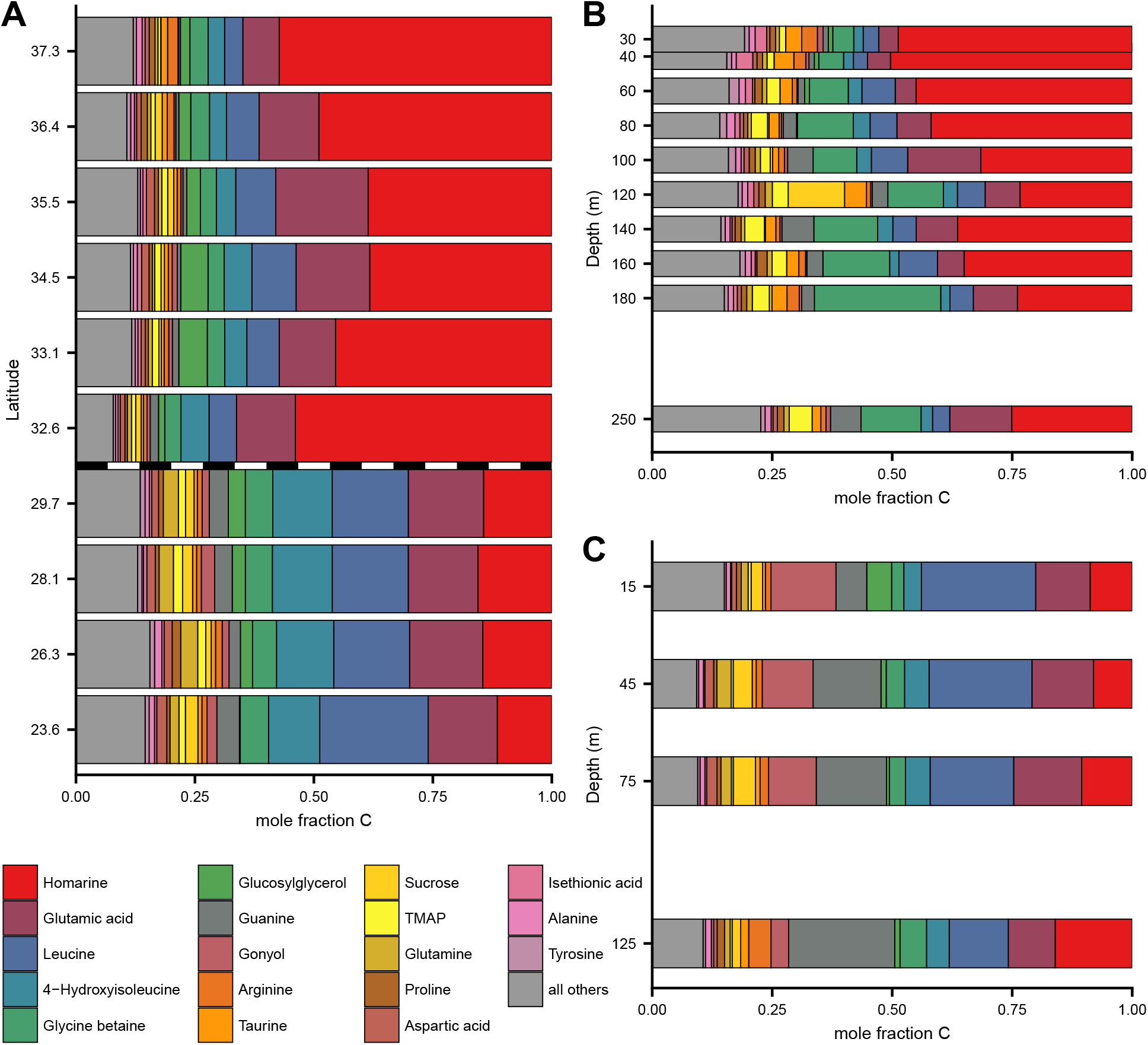
Most abundant 18 metabolites in environmental samples, presented as mole fraction of carbon of total identified metabolites. Meridional transect (A), with transition between NPSG to NPTZ shown as dotted line. Depth profile from NPTZ (B), and NPSG (C). Locations of samples are shown in Figure 1. These same data are presented as nmol C per L in Figures S8; full results in Table S10. Note that the y axis on (A) is not in latitudinal space for easier viewing and we have excluded DMSP from this analysis.

We quantified the same molecules in the 21 species of phytoplankton and one species of Thaumarchaeota (Table S3, Figures 5, S9). Most of the abundant compounds in the environment were also found in high abundance in at least some of our cultures, though many of the most abundant compounds were not ubiquitously observed across the cultures (e.g., glycine betaine and sucrose; Figure S9). We estimated the contribution of each metabolite to the carbon pool within each organism and compared this value to the surface samples of particulate metabolites in the field (Figure S5). This comparison yielded a consistency suggesting most of the surface particles contain compounds that have not been heavily reworked, corroborating previous work looking at macromolecule pools (37), particularly in the compounds found within the ‘core metabolome’ meta-cluster in Figure 3 (Figure S5). Comparing our environmental data sets to the culture data sets highlights compounds that were over-represented in either the culture data sets or environmental data set in a quantitative sense. For example, common compounds guanine and creatine and less well studied compounds like isethionic acid and dimethylsulfonioacetate (DMS-Ac), were all higher on a per carbon basis in the environment than in any of our cultures.

**FIG 5.**
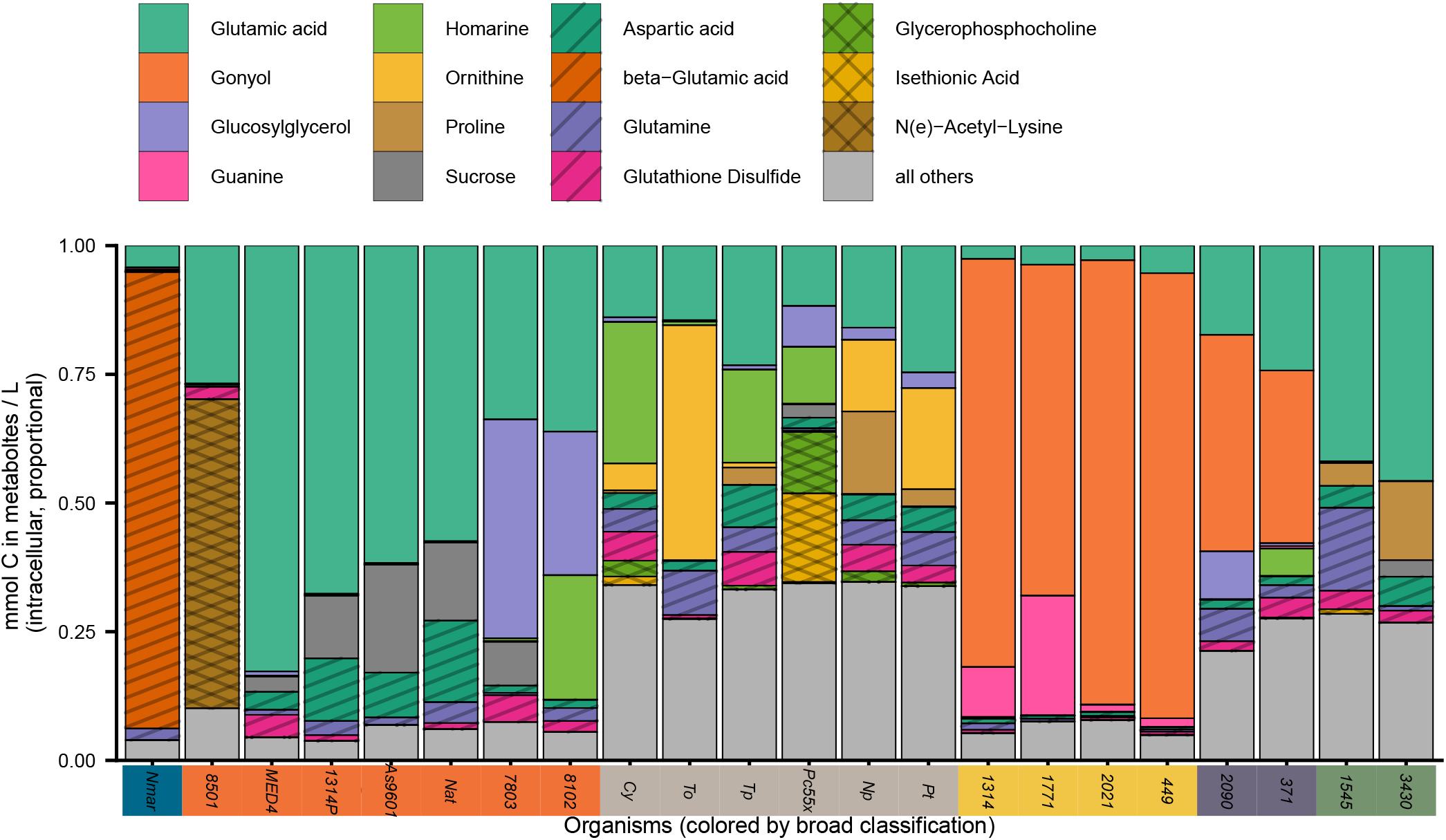
Identified metabolites in culture samples, presented as mole fraction of carbon of total identified metabolites, with at least the most abundant two compounds highlighted; full results in Table S8. Organisms are colored by broad taxonomic classification (as in Figure 2, blue = archaea, orange = cyanobacteria, grey = diatoms, yellow = dinoflagellates, purple = haptophytes, green = prasinophytes) and full description of cultured organisms found in Tables S3 and S7, full data in Table S8. We have excluded DMSP from this analysis since we cannot accurately quantify it using our methodology. Patterns are for added clarity with differentiating compounds.

### Homarine, an understudied metabolite of high abundance

The metabolite homarine (*N*-methylpicolinic acid) was present at 0.6 to 67 nM in marine particles, represented up to 3% of the total PC pool in our transect samples, and was the most abundant compound measured in our data sets (Figures 4, 6, and S9; Table S10). We found these concentrations surprising both in their absolute abundance and when compared to other more commonly studied polar metabolites known to accumulate in marine phytoplankton. For example, other studies have shown that homarine in marine particles is less abundant than the compatible solute glycine betaine (GBT) (38, 12), contrasting our findings. Both homarine and GBT are zwitterionic nitrogenous betaines that likely serve (at least in part) as compatible solutes. We also detected trigonelline (*N*-methylnicotinic acid), an isomer of homarine, albeit at much lower concentrations (1-300 pM in transect samples; Figures 6 and S9; Table S10). To our knowledge, trigonelline has not been previously detected in any marine samples, though it has been highlighted as an important component of labile carbon in terrestrial ecosystems due to its accumulation in higher plants (39).

**FIG 6.**
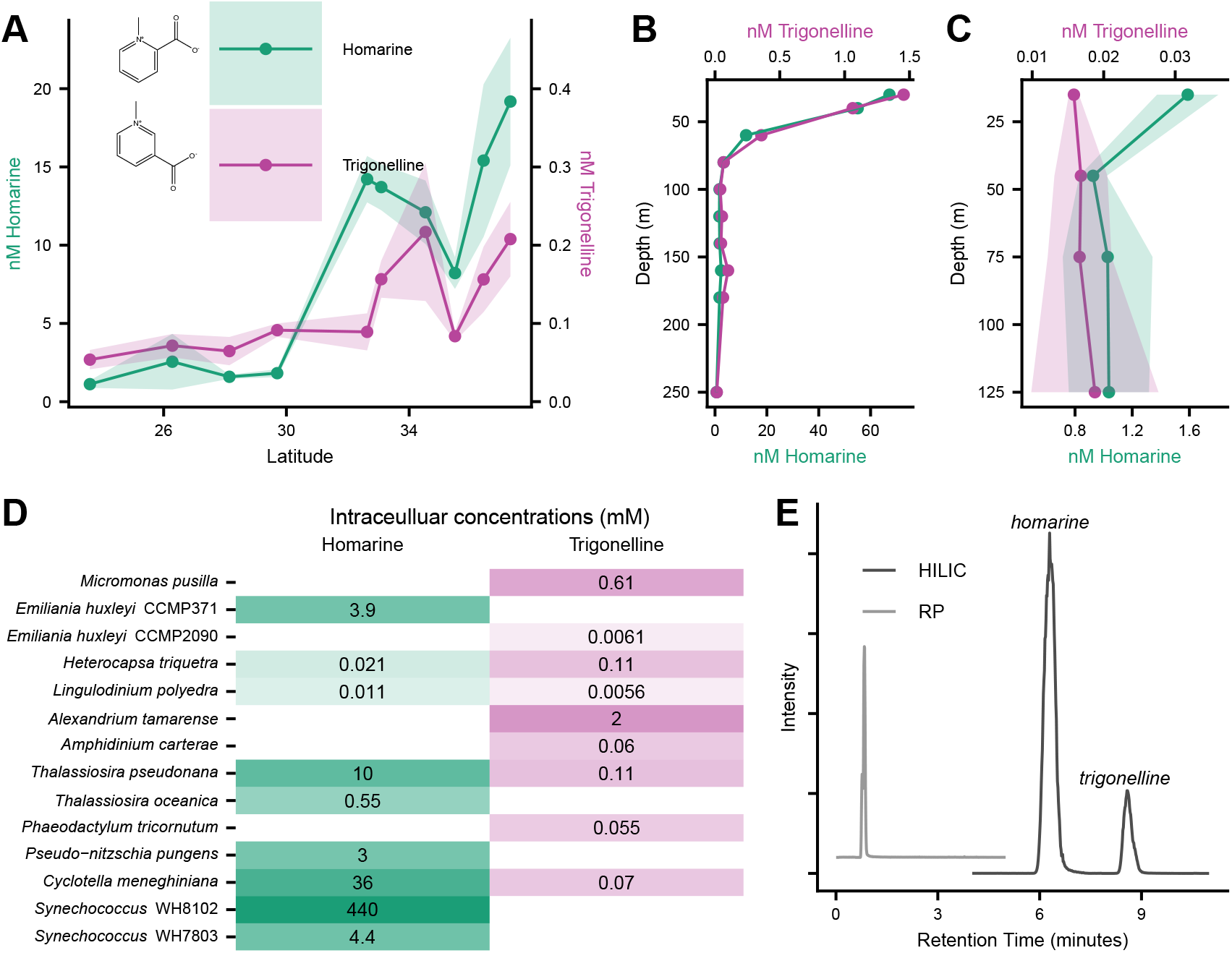
Homarine and trigonelline spatial patterns in meridional transect (A), NPTZ depth profile (B), NPSG depth profile (C). Intracellular concentrations of homarine and trigonelline in relevant organisms (D). Chromatograms show the separation under HILIC chromatography but not RP (E). Note the different scales for trigonelline and homarine (panels A-C) and depth (panels B and C).

In our cultured isolates, we detected homarine in both *Synechococcus* strains (intracellular concentration upto 400 mM), four of six surveyed diatoms (0.5-57 mM) and one strain of *Emiliania huxleyi*(a haptophyte, at 3.8 mM, Figure 6D, Table S8). Homarine has been observed in diatoms and *E. huxleyi* in previous studies (20,40,41,42,43,44), but has not been associated with the ubiquitous marine cyanobacteria *Synechococcus*. We estimated that homarine was 4.8% of the particulate carbon within *Synechococcus* strain WH8102. *Synechococcus* has been estimated to contribute 10-20% of global ocean net primary production at approximately 8 Gt C per year (45); by extrapolation this suggests up 0.5-1% global ocean net primary production could be attributed to this one molecule through *Synechococcus*, with potential for more production from diatoms. A caveat to this calculation is that homarine production is not quantitatively consistent among different strains of *Synechococcus; Synechococcus* WH7803 produced nearly 100 times less homarine (4-5 mM) under the same growing conditions (Figure 6, Table S8). This estimation is a first pass with the limited data at hand, and the sizable standing stock of homarine in our northern samples (about 2% of the PC) far exceeds what we would expect from the observed *Synechococcus* standing stock, which contributes less than 10% of the total PC pool (Figure S3). Homarine had a clear attenuation with depth in both of our depth profiles (within ‘mode *b*’ in the NPTZ and ‘mode *a*’ in the NPSG in Figure 1, shown in detail in Figure 6B, C). All together, these data support active production and cycling of this compound in the surface ocean that has been previously unnoticed.

Homarine showed a clear spatial pattern along our transect with nearly ten times higher abundance in the NPTZ (average 14.3 nM) than the NPSG (average 1.85 nM) (Figure 6A), which we hypothesize is a result of the changing phytoplankton community and increasing prevalence of *Synechococcus* around 32°N (Figure S3) and diatoms further north (indicated by increasing fucoxanthin around 34° N, Figure S6). Since *Synechococcus* standing stock cannot explain the observed homarine concentrations, we hypothesize that this compound may transfer to and accumulate in organisms beyond *Synechococcus*, which has been observed for osmolytes in other systems (44). Trigonolline followed a similar pattern along the transect, but with less pronounced increase in concentration from the NPSG (average 0.07 nM) to the NPTZ (average 0.14 nM) that was shifted more northward (Figure 6A). Homarine decreased sharply with depth in both depth profiles while trigonelline did not show appreciable attenuation in the NPSG profile (Figure 6B, C).

Biochemically, the sources and sinks for homarine and trigonelline are likely distinct. Trigonelline is produced from nicotinic acid (46), while homarine is decarboxylated from quinolinic acid, which is produced from tryptophan (47), though the exact enzyme that performs the decarboxylation has not been characterized. The first step of bacterial trigonelline degradation is the opening of the aromatic ring by the TgnA/TgnB oxygenase system (39). This enzymatic machinery is unlikely to operate on homarine due to steric hindrance in the ring-opening step. Supporting the differential catabolism of homarine and trigonelline, we saw that the model marine heterotrophic bacteria *Ruegeria pomeroyi* DSS-3 was not able to grow on homarine as effectively as trigonelline (Figure S10). Without characterized biosynthetic or degradation pathways for homarine, it is not surprising that this metabolite has not been identified as an important component of the labile organic carbon and nitrogen pools using gene-based techniques. Our spatial patterns and divergent observations of these compounds in our cultured organisms (Figure 6D) support distinct biological sources for these structurally similar compounds, demonstrating the intricate networks that exist in microbial communities rooted in the substrate-matched metabolisms.

Our observations of trigonelline and homarine were possible because of the chromatography methodology we employed (hydrophilic interaction liquid chromatography, HILIC) — the compounds would not be resolved in time in more commonly employed reversed phase chromatography (23, 25) due to their high polarity and same empirical formula (and therefore exact mass, Figure 6E). This is also true for many sets of isomers of known compounds (e.g., sarcosine, *β*-alanine, and alanine; homoserine and threonine; *β*-glutamic acid and glutamic acid) as well as unknowns (e.g., inosine and another unidentified metabolite with the same *m/z;* two metabolites with an *m/z* of 236.1492; see Table S5). We bring attention to this detail to highlight the power of incorporating cutting-edge analytical capabilities to study microbial ecology — without HILIC chromatography, we would not have been able to accurately measure many of the most abundant polar compounds.

### Organic sulfur compounds

Six of the top 30 most abundant compounds in our environmental samples were organic sulfur compounds. These compounds fall into two general categories; sulfoniums ([SR_3_]^+^) and sulfonates ([RSO_3_]^-^). We detected the well-studied sulfonium compound DMSP, though our methods likely underestimated the concentration due to compound instability in methanol-based extractions (48). Using our untargeted approach, we putatively identified two additional sulfonium compounds, dimethylsulfonioacetate (DMS-Ac) and 3-5-dimethylsulfonio-3-hydroxypentanoate (gonyol), as prominent peaks in our environmental samples. We later obtained standards that confirmed these identifications and enabled quantification that revealed gonyol as among our most abundant compounds with a particularly high concentration (up to 2.5 nM) in the NPSG depth profile (Figure S11). Gonyol was named after the dinoflagellate *Gonyaulax polyedra* (49), and gonyol was present in high concentration in all four dinoflagellate strains (81-196 mM) and in lower concentrations in the haptophytes (23-61 mM, Figures S9, S11, Table S6). The environmental samples contained more DMS-Ac per unit carbon than culture samples, suggesting a source of this compound in the environment not reflected in the cultured phytoplankton (Figure S5). Both of these compounds share structural similarity with DMSP, and disrupt bacterial DMSP degradation pathways (50). Thus, predicting marine DMS production from DMSP may be complicated by these highly abundant compounds. Although marine organic sulfur has gained much attention with regards to its massive inventory (51) and role in microbial processes (52), ours are the first observations of these understudied sulfoniums in natural marine systems.

### Remaining unidentified compounds

Many of the metabolites with interesting patterns across space and taxonomy could not be identified (Table S5). For example, the mass feature “I121.0684R10.7” has a *m/z* of 121.0684 and major peak in its MS^2^ fragmentation spectra of *m/z* 63.02703 (Table S5). This metabolite likely has the empirical formula of C_5_H_12_OS and was observed in 19 of the 21 phytoplankton species, attenuated with depth, and had a distinct maximum from 32-34°N in the meridional transect (Table S5). Unfortunately, none of the possible matches to these compounds have fragmentation data in the major mass spectral databases; without an identification, we cannot quantify this compound. It is very likely that within these unidentified compounds are more underappreciated compounds involved in the microbial loop — a fruitful endeavour for future oceanographers, mass spectrometrists, and biochemists alike.

### Conclusions

Small molecules within marine particulate organic matter contribute to the dissolved organic matter pool after excretion, cell lysis, or sloppy feeding. Once in a dissolved form, other organisms in the environment may be able to use these compounds as substrates as sources for carbon, nutrients, and energy (53,54), if they have the required enzymatic machinery to access these resources. These small molecules may also act as chemical attractants or deterrents for organisms and therefore assist in shaping microbial communities. By directly observing small molecules in both field particulate material and cultured phytoplankton, we show that small molecules in natural marine systems are determined in part by the taxonomy of the phytoplankton community. This suggests that to access these pools of labile organic carbon the wider microbial community must be adapted to phytoplankton community composition. By quantitatively contextualizing our metabolomics data sets, we uncover a rich set of compounds that likely fuel the microbial loop that have been previously overlooked. Cycling of organic matter thus depends both on the amount of primary productivity and phytoplankton composition — *who* matters on a chemical level.

## MATERIALS AND METHODS

### Environmental sample collection

Samples were collected for environmental metabolomics of particulate material at locations shown in Figure 1. Samples for the NPSG depth profile were collected aboard the R.V. Kilo Moana cruise KM1513 on July 31,2015 from four depths (15, 45, 75, and 125 m); we reported on these samples in a previous publication (16). Samples for the meridional transect were collected on cruise KOK1606 aboard the R.V. Ka’imikai-O-Kanoloa from April 20 to May 2, 2016, all at approximately 15 m. Samples for the NPTZ depth profile were collected during MGL1704 aboard the R.V. Marcus Langseth at seven depths between 30 and 250 m on June 3, 2017. At each sampling location and depth, single, duplicate, or triplicate filters were collected for environmental metabolomics, as previously described (16), using either niskin bottles or the uncontaminated underway seawater intake. Table S1 has summarized descriptions of the samples collected for metabolomics, with full description of each sample (including time of collection) in Table S4. In short, samples (4-15 L each) were collected into polycarbonate carboys, filtered onto 147 mm 0.2μm PTFE filters using a peristaltic pump, flash frozen in liquid N_2_, and stored at −80°C until extraction. In addition to our samples, we filtered duplicates of methodological blanks by filtering seawater through two 0.2 *μ*m PTFE filters in series and used the second filter as the blank. This blank is especially important to parse metabolite signals from contaminants as well as compounds within the residual dissolved pool and salt matrix adsorbed during filtration.

### Pure cultures and sampling

In addition to environmental samples, we analyzed metabolomes of cultured representatives of marine phytoplankton that were grown and analyzed on the same LC/MS system as previously presented (5). Media, light, and temperature were chosen for optimal growth of each species and are reported in (5). In short, axenic phytoplankton were cultured in controlled laboratory settings and harvested under exponential growth using a gentle vacuum filtration onto 47 mm Durapore filters (pore size 0.2 *μ*m). Samples were flash frozen in liquid N_2_ and stored at −80°C until extraction. In addition to samples, media blanks corresponding to each media type were harvested and served as matrix blank to each corresponding phytoplankton sample. In order to estimate intracellular concentrations of metabolites, we used biovolume estimates from (5).

We also grew *Nitrosopumilus maritimus* strain SCM1 and harvested under exponential growth. Pure culture of Marine Group I Thaumarchaeota *Nitrosopumilus maritimus* strain SCM1 was maintained in liquid mineral medium with 1 mM ammonia (55) at 30°C in the dark without shaking. The growth of *N. maritimus* was monitored by measuring nitrite production and cell abundance. Nitrite concentration was determined spectrophotometrically using the Griess reagent (56). Cell counts were determined using Moviol-SYBR Green I staining protocol as previously reported (57) with a Zeiss epifluorescence microscope to count 15 random fields of view for each sample with 30 to 200 cells per field. Mid-exponential phase cells were harvested using a gentle vacuum filtration on 0.22 *μ*m Durapore membrane filters (Millipore Co., MA, U.S.) and stored at −80°C until metabolite extractions. These archaea have a biovolume of approximately 0.023 *μ*m^3^ (58).

We estimated carbon contents for all the cultures from cellular volume (59), using an empirical relationship between flow cytometry-based cell size and PC (60), or using previous direct measurements (60,61,62). An abbreviated sample description is given in Table S3; full sample descriptions are in Table S7 (including carbon estimates and the method used for each species).

### Additional oceanographic data

Samples for particulate carbon were sampled and processed as in (63). Underway flow cytometry data were acquired and processed as in (60). Samples for pigment analysis were filtered onto GF/F filters (Whatman), stored in snap-cap tubes, wrapped in aluminum foil, and flash-frozen. Samples were analyzed for high performance liquid chromatography (HPLC)-based measurements of total chlorophyll (monovinyl + divinyl), fucoxanthin and other photosynthetic and photoprotective pigments. These analyses were made in the Oregon State University HPLC facility via a Waters 996 absorbance photodiode array detector in combination with a Waters 2475 fluorescence detector according to the protocol of (64).

### Homarine bioavailability experiment

To test if homarine was as bioavailable as trigonelline in marine systems, we cultured the model marine heterotrophic bacteria *Ruegeria pomeroyi* DSS-3 under different primary carbon sources and observed its cell density. DSS-3 were streaked to isolation on 1/2 ytss agar plates (1.25g tryptone, 2g yeast extract, 10g sea salts, 8g agar per 500 mL MQ water) from frozen glycerol stocks at room temperature 3 d. A single colony was inoculated into artificial L1-bac seawater media (described below) supplemented with acetate (final concentration of 50 nM). This culture was grown overnight at room temperature at 200 r.p.m at 30°C. Next, a 96 well plate was prepared with 90 *μ*L of fresh media described above (without additional carbon) in all wells. In 8 wells, we added 5 *μ*L of the overnight inoculum and 10 *μ*L of water (no additional carbon treatment). In 8 wells, we added 15 *μ*L of water and no inoculumn (negative control). In the remaining wells, we added 5 *μ*L of the overnight inoculum and 10 *μ*L of either acetate, homarine, or trigonelline (all at 100 nM carbon, *n* = 8 for each, acetate serving as positive control). Plates were covered in a breathable sealing membrane (Breathe-Easy) and placed into a platereader (Biotek Synergy H1MF). Cultures were grown at 30°C, shaken every 2 minutes for 3 seconds, and monitored via absorbance at 600 nm every 2 minutes (immediately after shaking).

Artificial L1-bac seawater media was prepared using MQ water with 28 g Sigma sea salts, trace and marcro nutrients based on the recipe from the National Center for Marine Algae and Microbiota (without silica), nitrogen, and vitamins as in (18), Sigma M5550 MEM essential amino acids (1:1000 dilution), Sigma M7145 MEM non-essential amino acids (1:2000 dilution). Salt water was autoclaved in combusted borosilicate glass containers and all additions were made from filter sterilized stocks. Final media was filter sterilized using a 0.22 *μ*m PVDF membrane bottle top filter.

### Metabolite data acquisition

Metabolites were extracted as previously described (16). Briefly, filters were bead-beaten three times in 30 s bursts over 30 minutes (kept at −20°C between bursts) in 1:1:2 methanol, water, dichloromethane and separated into two fractions: a polar aqueous extract (methanol and water extractable) and an organic extract (dichloromethane extractable). We used the same internal standard suite at the same injection concentrations as in Boysen *et al*. (16) to train normalization and monitor instrument stability. After drying under clean N_2_ all samples were reconstituted in 400 *μ*L water.

The polar fraction of this extract was analyzed on both reversed phase (RP) and hydrophilic interaction chromatography(HILIC) using the same solvents, columns, and gradients as previously reported (16). We diluted the KOK1606 samples (1 part sample to 2 parts water) and MGL1704 samples (1 part sample to 1 part water) which helped with signal stability over the course of the runs. Internal standards were added during the dilution step and were the same concentrations in all analyzed samples to aid in quantitative comparisons between sample sets. We injected 2 *μ*L of sample onto the column for HILIC analysis, and 5 *μ*L (for environmental samples) or 15 *μ*L (for culture samples) for RP analysis.

Both LC configurations (RP and HILIC) were analyzed on a Thermo Q-Exactive (QE) mass spectrometry in full scan mode for quantitative data, or data dependent acquisition (DDA) for fragmentation. Full scan analyses were conducted as in Boysen *et al*. (16); pooled samples were run in DDA mode for MS^2^ fragmentation as described in Heal *et al*. (20).

### Metabolomic data processing

To compare our field data with our culture data, we used our untargeted data from the meridional transect (36 surface samples) as our template to examine the other sample sets. To do this, we used an established untargeted metabolomics approach (detailed below) to acquire a list of curated, dereplicated, and high-quality mass features. With this curated list, we then searched for the same mass features in the remaining field and culture sample sets. This allowed us to compare relative abundances of these mass features within each sample set, with high confidence in the shared identity of these compounds between sample sets.

Untargeted metabolomics data from transect samples were converted with MSConvert (65) and processed through XMCS (66, 67, 68), using the same parameters for XCMS and methodological blank filtering as previously reported (20). Next, we normalized for obscuring variation (non-biological variability inherent to LC-MS analysis) using B-MIS normalization (16). Like in Heal *et al*. (2019), we disregarded peaks that did not demonstrate acceptable replicability in the pooled samples (coefficient of variance > 30%); we also removed peaks that showed greater average variability between biological replicates than over the whole sample set as in previous work (4).

In untargeted metabolomics, multiple mass features can correspond to one metabolite due to natural abundance isotopes, adducts, or multiply charged ions. Like in Heal *et al*. (2019), to avoid putting extra statistical weight on these isotopes and adducts, we identified mass features that were likely ^13^C, ^15^N, or ^34^S isotopologues of other mass features. We extended this search to include adducts of Na^+^, NH_4_^+^, K^+^, (for positive ionization) and Cl^-^ (for negative ionization), as well as for doubly charged ions of mass features whose M+H ion was present. We performed these searches within each three second (for RP) or six second (for HILIC) corrected retention time window and discarded these mass features from downstream statistical analyses.

For the largest 200 peaks in our HILIC analysis (positive and negative analyzed separately) and RP analysis, we exported the *m/z* and retention time information to Skyline (69) for closer inspection. XCMS peak picking algorithms assume a normal Gaussian shape for peaks (66, 67, 68), which often results in poor integrations for compounds that do not achieve this shape during chromatographic separation; these peaks are often removed during our CV filter or manual peak quality verification. Therefore, we also imported a list of compounds we regularly target (see (16) for full list of standards) and manually integrated these compounds in each of the samples (first removing compounds that were picked during the peak picking step). In Skyline (69), we integrated these peaks (both the untargeted and known compounds) in the transect data set since XCMS often results imperfect integrations that can introduce non-biological variability to metabolite abundances (70). Next, we eliminated mass features that were not present in at least 50% of the transect samples and also removed peaks that were not (on average) three times larger than the matrix blank. These stringent filters in the transect data set allowed us to use a culled number of high-quality mass features that are common in surface seawater particles as a fingerprint of metabolite pools. In all, we obtained 149, 74, and 90 high-quality, manually integrated peaks in HILIC-positive ionization (HILICPos), HILIC-negative ionization (HILICNeg), and RP, respectively. For these quality mass features, we searched for corresponding MS^2^ scans in the data dependent acquisition (DDA) files and applied a filter to remove low abundance fragments in the exact manner as reported in (20).

With this list of high-quality mass features (referred to in the text as metabolites), we extracted the exact masses and integrated peaks at the same retention times in the two other environmental data sets (NPSG and NPTZ depth profiles) and the culture data sets. We also integrated our internal standards (in exact concentrations as in (16)) and performed B-MIS normalization (16) across our environmental data sets which minimizes the variability present during analysis (not biological variability). This resulted in *adjusted areas* of each compound in each sample that are quantitatively comparable within each sample set (but not between). Since the phytoplankton data sets are not in a consistent matrix and were analyzed in several different batches, we did not attempt to use B-MIS to normalize across the organisms. Instead, we kept the raw peak area, normalized it to the biovolume analyzed, and made semi-quantitative comparisons on the log_10_ transformed biovolume-normalized peak areas. The log-io transformations ensures that only large differences are evaluated as contributing to variability between samples, well beyond matrix variability or instrument performance.

As in (20), we used the ranking system outlined in (71), to attempt to identify the quality mass features present in these sample sets in an automated fashion. We searched an internal database of compounds with known exact *m/z* and retention time on the LC-MS configurations used in the lab (found at https://github.com/Ingalls-LabUW/Ingalls/Standards), publicly available MS/MS^2^ spectral databases (72, 73, 74, 75, 76), and to compounds in the KEGG database (77, 78) (based only on *m/z*).

### Calculating concentrations

Commercially available standards were analyzed in the same batch as each of the three environmental data sets. For this subset of com-pounds, we calculated absolute concentrations, similar to previous work (21, 40, 12). In short, we applied the following calculation for each analyte.

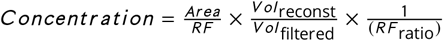

Where *RF* is the response factor 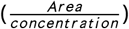 of each compound at known concentration in water. Standards are run before and after each run on each instrument, therefore, an RF for each compound is obtained within each batch. *Vol*_reconst_ is the volume that the samples were reconstituted into; *Vol*_filtered_ is the volume filtered in the field (for environmental samples) or the total estimated biovolume collected (for culture samples); *RF*_ratio_ is the 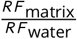 these compounds in a matrix of marine particulates (as described in Boysen *et al*. (16)), we calculated *RF*_ratio_ using samples from the transect data set which were applied throughout. We calculated a *RF*_ratio_ as in (16) using a representative environmental matrix sample. Values for *Vol*_filtered_ for each sample are reported in Tables S4 and S7; *Vol*_reconst_ was 400 *μ*L for each sample.

Several compounds were identified in the transect data set and purchased and analyzed using the same LC/MS method at a later date, which we quantified using the same approach as in (40). Because the *RF* for each compound can vary substantially between analytical runs, we used a relative response factor (*RF*_relative_) to estimate RF and calculate the concentrations of these compounds in earlier runs. To calculate *RF*_relative_, we matched compounds with a standard that had been analyzed in all sample sets that share the same column, ionization state, and some structural similarity (matched standard). For instance, for the compound DMS-Ac, we matched it to another dimethylated sulfonium zwitterion, DMSP. After the samples were analyzed, we analyzed these new standards and the other standards on the same LC-MS set up as our sample set and calculated *RF*_relative_ using the following formula:

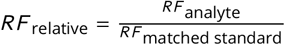

Then we used this *RF*_relative_ and the *RF*_matched standard_ to calculate the concentration of the analyte from earlier runs. For a full explanation for how each compound was quantified in each sample set and for the matched standard used for each compound (when necessary), see Table S11.

### Statistical approaches

For multivariate statistics on environmental samples, peak areas (adjusted via B-MIS for instrumental variability and normalized to water volume filtered) were standardized to the total peak area observed for each mass feature across each sample set. For each mass feature in the cultured organisms, log_10_ transformed peak areas were standardized to the maximum log_10_ peak areas observed across all cultured organisms. We used two different multidimensional approaches on these data sets, both non-metric to accommodate for the high variable to sample ratio and non-normal distribution of peak areas in our data sets. This prevents overfitting which can be a problem in other multidimensional approaches in metabolomics (79). We used a non-metric dimensional scaling (NMDS) analysis (80) based on a Euclidean distance matrix of standardized peak areas to visualize overall metabolic differences between samples along our transect samples. We assessed dimensionality of the NMDS by examining a scree plot, calculated the probability with a Monte Carlo permutation which resulted in a low stress ordination. We accompanied this with an analysis of similarities (ANOSIMS) (81) to discern differences between the oceanographic regimes we sampled as well as time of sampling. Data transformation, standardization, NMDS, and ANOSIMS statistics were performed in R using the vegdist (v2.4-2) or vegan (v2.4-2) packages.

Next, we employed a *k*-medoids based clustering approach (82) which aggregates metabolites based on patterns across samples. We performed this clustering on the combined culture data sets and on the three environmental sample sets separately (four total *k*-medoids analyses) using the clara function in the cluster package (v2.1.0) in R. This non-supervised clustering technique is exclusive and non-hierarchical which assigns each mass feature into one cluster, or mode. The metabolites within each mode have similar patterns of abundance across samples. We chose the appropriate number of modes for each sample set by selecting the mode number that resulted in a local maximum average silhouette width between samples.

Finally, we investigated whether the resulting modes of metabolites from the four separate *k*-medoids analyses shared metabolites beyond a random assignment. Essentially, we asked if the patterns in the 2016 transect samples could be explained in part by patterns in metabolites across the available culture data or could be recapitulated in the depth profile sample sets. To test for over-represented sharing of metabolites between modes, we used a Monte Carlo resampling technique to simulate the random frequency of shared metabolites using 1000 permutations. We then compared the observed frequency of shared metabolites to the permutations to estimate the *p*-value of our observed shared metabolites.

For all data analysis, we used R v4.0.0. Codes for figures, tables, and data analysis are found at https://github.com/kheal/Gradients1_SemiTargeted3. Raw data for metabolomics samples are deposited at Metabolomics Workbench; all cultures are under project ID is ST001514, project IDs for environmental samples listed in Table S4.

## SUPPLEMENTAL MATERIAL

Supplemental Figures S1–S10 and Tables S1–S3 supplied in a combined document following the references with more detailed legends. Supplemental Tables S4-S11 are supplied as a combined .xlsx, full legends supplied here.

**FIG S1**. Particulate carbon over the April 2016 transect and total quantifiable particulate metabolites.

**FIG S2**. Nonmetric multidimensional scaling comparison of metabolite profiles on the KOK1606 transect.

**FIG S3**. Populations of picoeukaryotes and picocyanobacteria over the transect, as observed via underway flow cytometry (SeaFlow).

**FIG S4**. Nonmetric multidimensional scaling comparison of metabolite profiles among the cultured organisms.

**FIG S5**. All identified and quantified compounds compared in carbon space in cultures and environmental samples.

**FIG S6**. Concentration of the pigment fucoxanthin over latitude in transect samples.

**FIG S7**. *β*-glutamic acid and arsenobetaine depth profiles from the NPTZ.

**FIG S8**. Most abundant metabolites in environmental samples, presented as nmole carbon per L.

**FIG S9**. All identified compounds, with average and standard deviation of observed concentrations in surface seawater particles or in culture samples.

**FIG S10**. Growth curves of *Ruegeria pomeroyi* DSS-3 with three different carbon sources: acetate (positive control), homarine, and trigonelline, and no additional carbon (negative control).

**TAB S1**. Summary of samples collected and analyzed in this study.

**TAB S2**. Summary of physical and chemical parameters on April 2016 cruise. Reprinted with permission from (27).

**TAB S3**. Summary of cultured organisms analyzed in this study.

**TAB S4**. Full sample descriptions for environmental samples. Binned latitude is the latitude used for plotting when aggregating the data. This table is supplied in a .xlsx file.

**TAB S5**. Peak areas from environmental and culture samples. MassFeature-Column is the identifier for each distinct mass feature (identified or not), as in all tables. Identification is the best identification we have for the mass feature; Confidence is based on (71), with 1 as unequivocal (and quantifiable); mz is mass to charge ratio observed; rt is retention time (in seconds); Column is the chromatography method used (HILIC or RP); z is charge state in which the mass feature was observed (1 is positive, −1 is negative). This table is supplied in a .xlsx file.

**TAB S6**. Assignments of mass features compounds to modes and metaclusters (if applicable) as a result of the *k*-medoids clustering summarized in Figures 1, 2, and 3. MassFeature_Column is the identifier for each distinct mass feature (identified or not), as in all tables. Identification is the best identification we have for the mass feature; Confidence is based on (71), with 1 as unequivocal (and quantifiable); mz is mass to charge ratio observed; rt is retention time (in seconds); Column is the chromatography method used (HILIC or RP); z is charge state in which the mass feature was observed (1 is positive, −1 is negative). MS2 is the MS^2^ spectra for the peak in a pooled sample from the transect sample set. This table is supplied in a .xlsx file.

**TAB S7**. Full sample descriptions for culture samples. All volumes in *μ*L. Carbon estimates are based on (59, 60, 61, 62), as indicated. This table is supplied in a .xlsx file.

**TAB S8**. Quantified metabolites from all culture samples. Compound names are given as reported in figures and as more complete names for clarity. All values intracellular in *μ*mol metabolite per L. Sample identifiers are elaborated in Table S7. This table is supplied in a .xlsx file.

**TAB S9**. Total quantifiable metabolites as a fraction of the particulate carbon and nitrogen pools. All measurements are mean (standard deviation), except when *n* = 1. Standard deviations of calculations (percentages) are propagated. Note that we do not have bulk particulate carbon or nitrogen measurements paired with the depth profiles. This table is supplied in a .xlsx file.

**TAB S10**. Quantified metabolites from all environmental samples. Compound names are given as reported in figures and as more complete names for clarity. All values areμmol metabolite per L seawater. Sample identifiers are elaborated in Table S4. This table is supplied in a .xlsx file.

**TAB S11**. Quantification method for each quantified metabolite in each sample set. Proxy compound (when applicable) is the compound by which a relative response factor (RF) was calculated. This table is supplied in a .xlsx file.

## ACKNOWLEDGMENTS

We would like to thank Grace Workman, Natalie Kellogg, Regina Lionheart, Rhonda Morales, and Alexa Wied for assistance in growing or enumerating cultures, extracting environmental metabolomics samples, and preliminary data processing. We acknowledge the science team and crew of KM1314, KOK1606, and MGL1704 cruises (particularly Rachelle Lim and Ryan Grossmann) for sample collection. We thank G.Pohnert for the generous donation of gonyol and DMS-acetate standards. Cultures were provided by D. Stahl, S. Chisholm, A. Coe, M. Moran and M. Saito. This work was supported by grants from the Simons Foundation (385428 to A.E.I; 426570 to E.V.A., A.E.I., A.E.W., and R.M.B.; 548565 to W.Q.; 598819 to K.R.H.), the National Science Foundation (NSF GRFP to A.K.B. and K.R.H.; NSF OCE-PRF 1521564 to B.P.D., NSF IGERT Program on Ocean Change to A.K.B).

## SUPPLEMENTAL FIGURES

**FIG S1.**
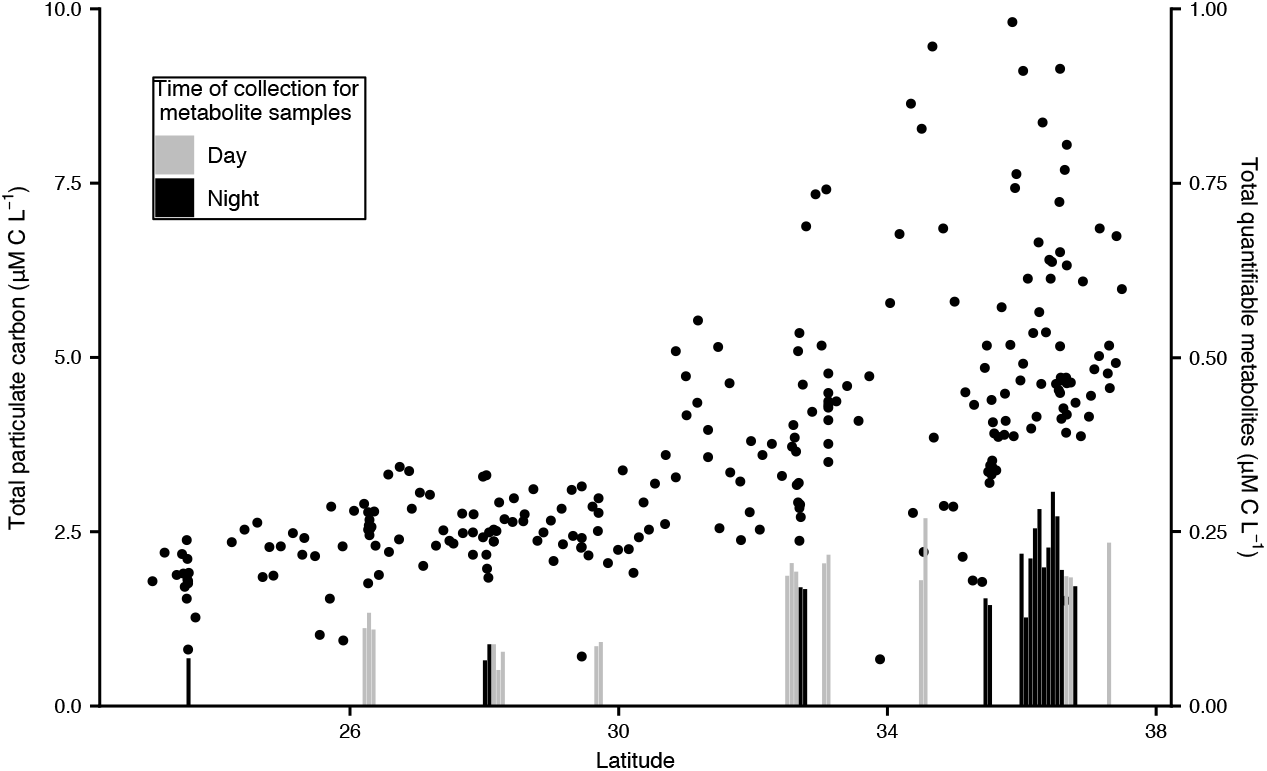
Particulate carbon over the April 2016 transect (dots) and total quantifiable particulate metabolites (bars). Note that the total quantifiable metabolites are scaled at *x* 10 for visibility and dodged in latitude to show each individual replicate.

**FIG S2.**
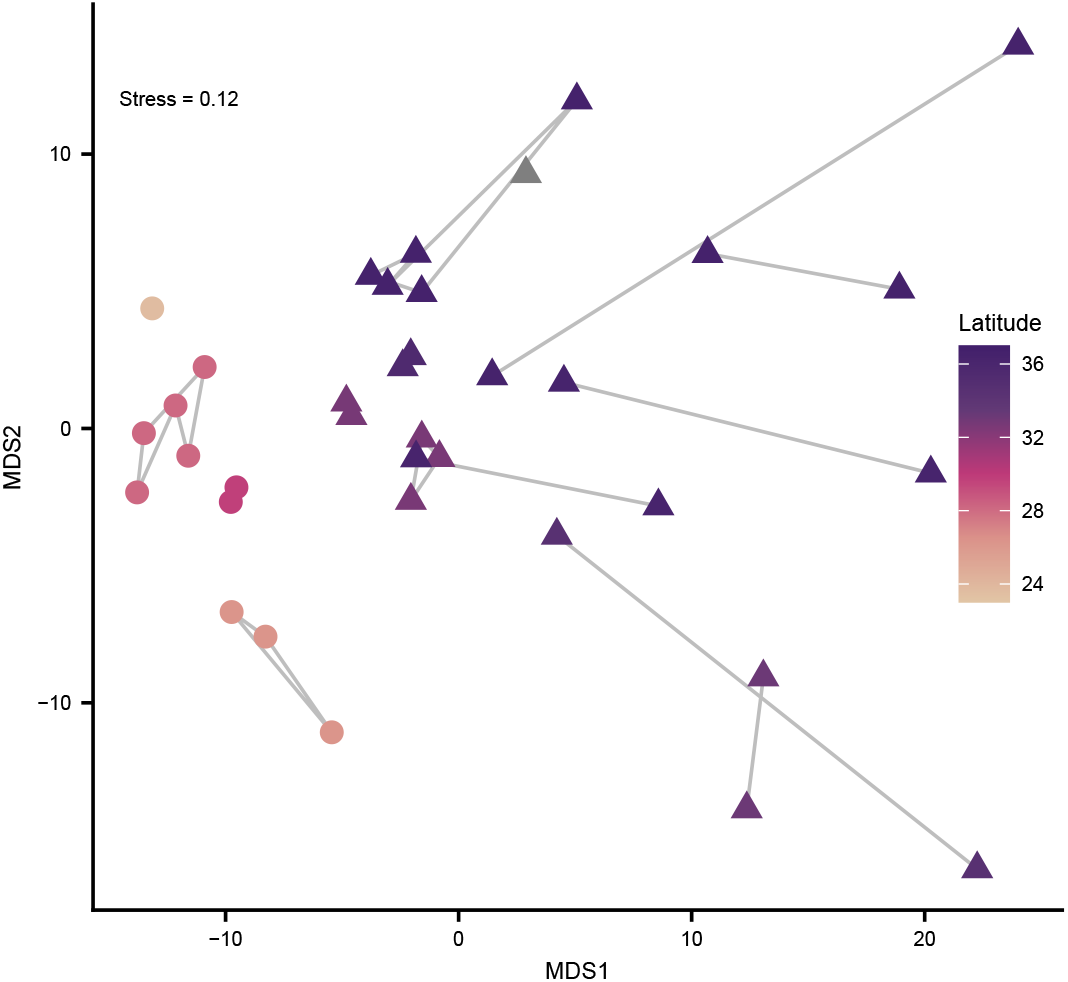
Nonmetric multidimensional scaling comparison of metabolite composition on the KOK1606 transect, *p* < 0.01 by Monte Carlo permutation. Colors are based on latitude of samples; biological replicates shown connected, with samples from NPSG in circles and NPTZ in triangles. This is based on euclidean distance between standardized adjusted peak areas.

**FIG S3.**
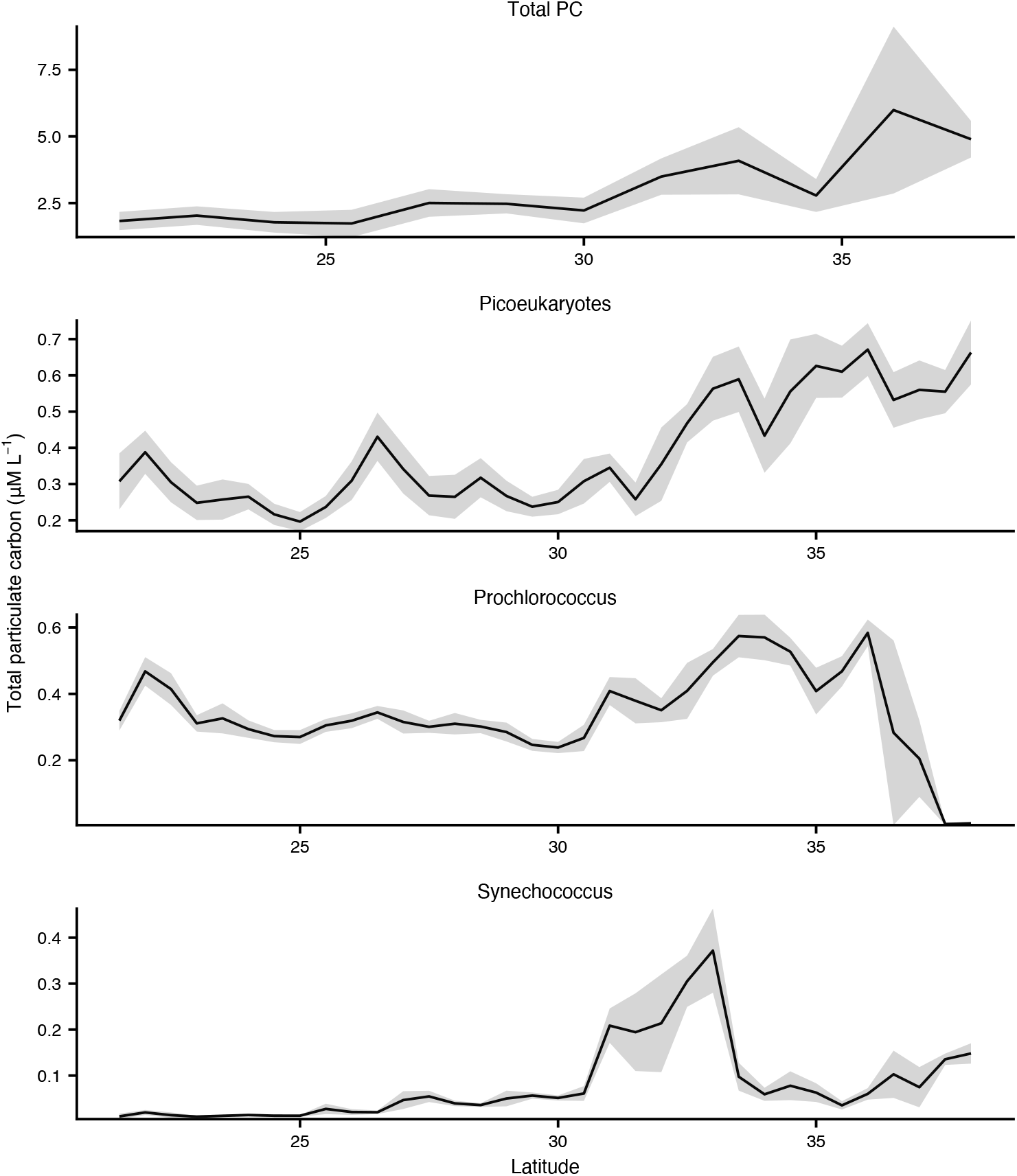
Particulate carbon (total, and estimated by particular populations of phytoplankton as observed via underway flow cytometry (SeaFlow). Phytoplankton carbon are binned every 0.5 degree latitude, total PC binned every 1 degree latitude. Grey shading shows standard deviation. This is the northbound transect only.

**FIG S4.**
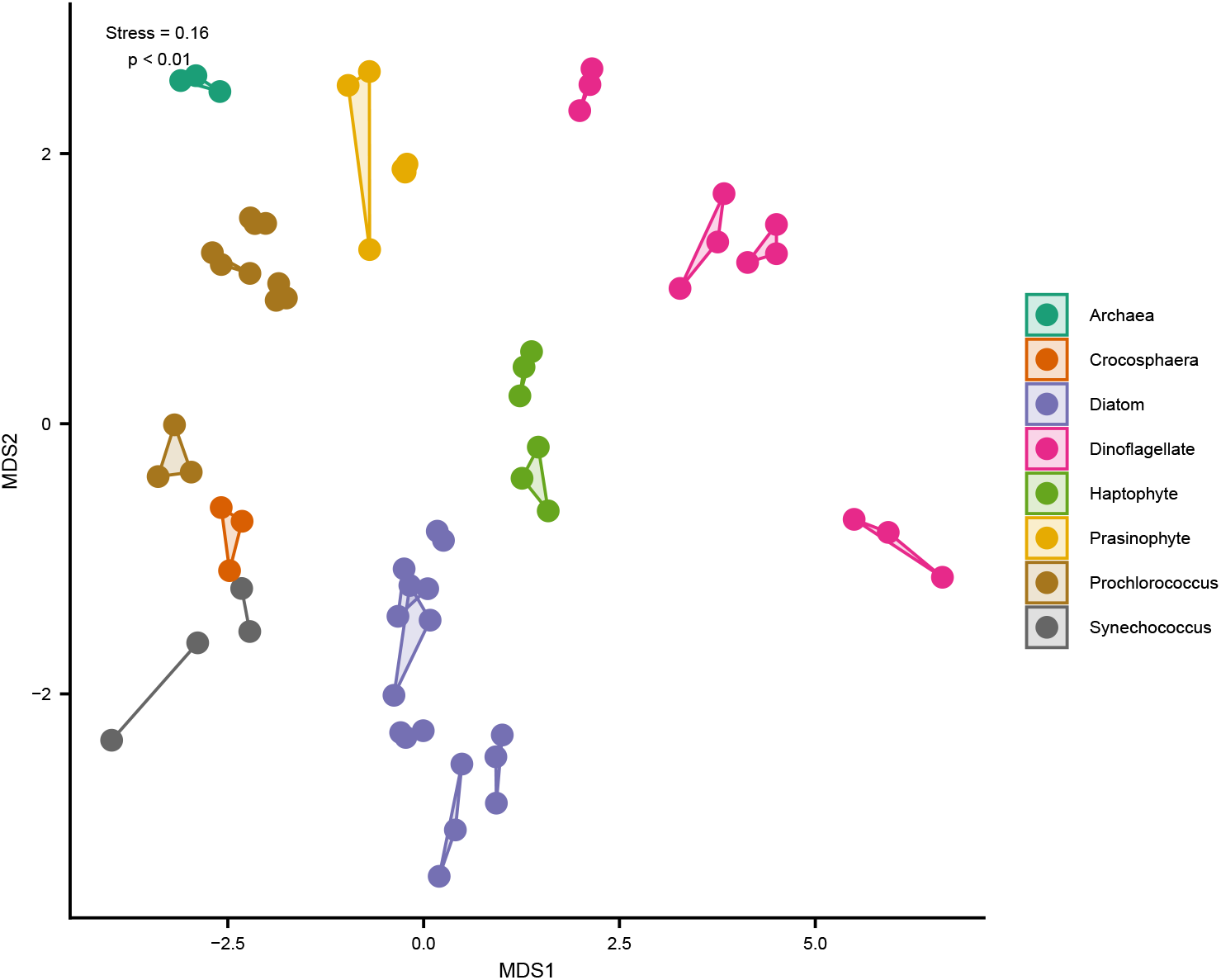
Nonmetric multidimensional scaling comparison of metabolite profiles among the cultured organisms, *p* < 0.01 by Monte Carlo permutation. Colors are based on broad taxonomic groups; biological replicates shown connected. This is based on euclidean distance between standardized adjusted peak areas.

**FIG S5.**
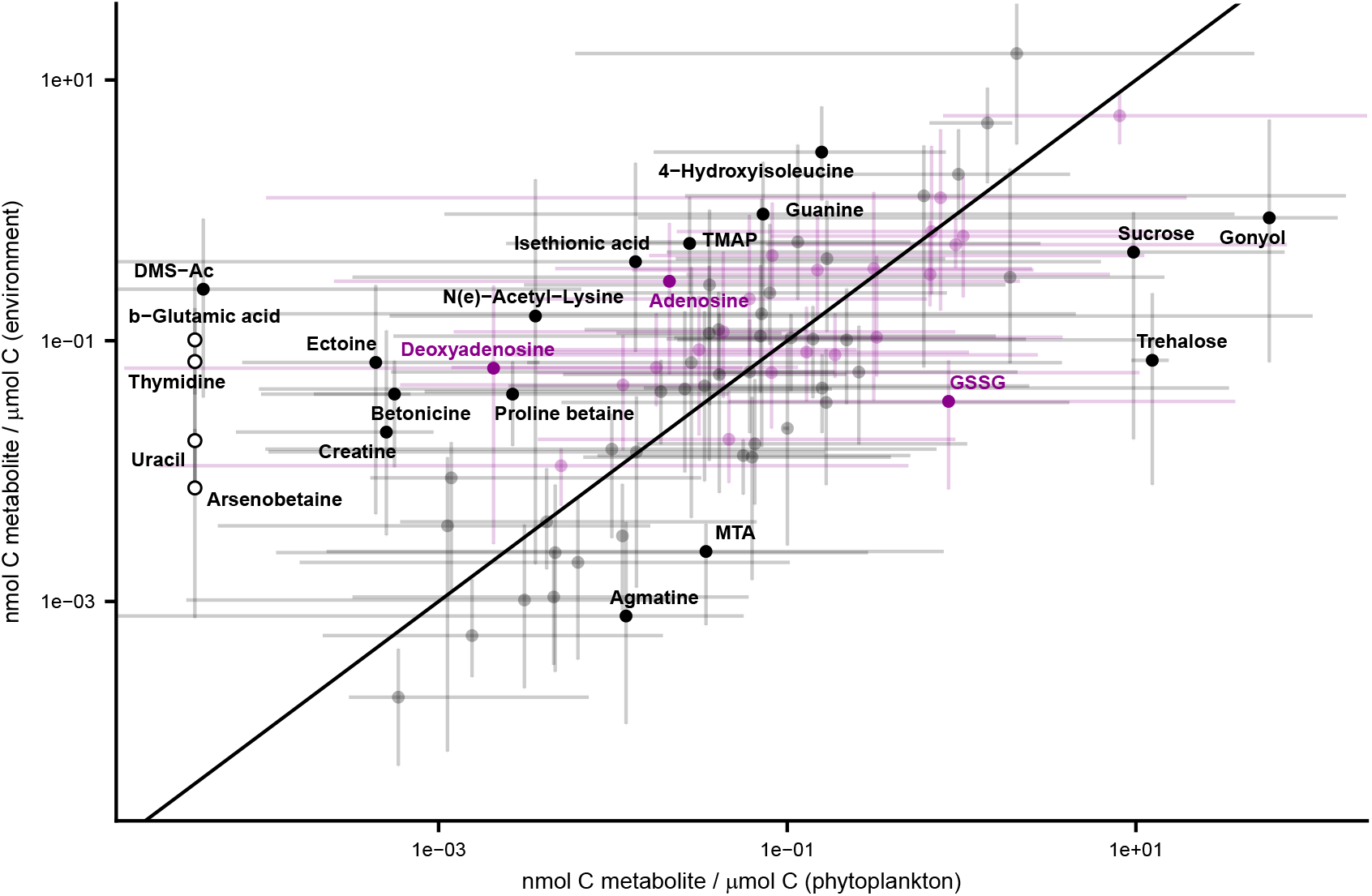
All identified and quantified compounds. Each compound is shown as a dot with error bars, representing median observation and range of observations, respectively (excluding instances where we did not observe the compound), with 1:1 line plotted. Compounds that were not detected in phytoplankton are shown as open circles (x value is arbitrary). Compounds assigned to the “core” meta-cluster in Figure 3 are plotted in purple. Compounds are highlighted when median observation are ten times higher in either environmental sample set or the culture sample set (in carbon space). These values are also reported in Table S10. Note that both axes are in log_10_ space.

**FIG S6.**
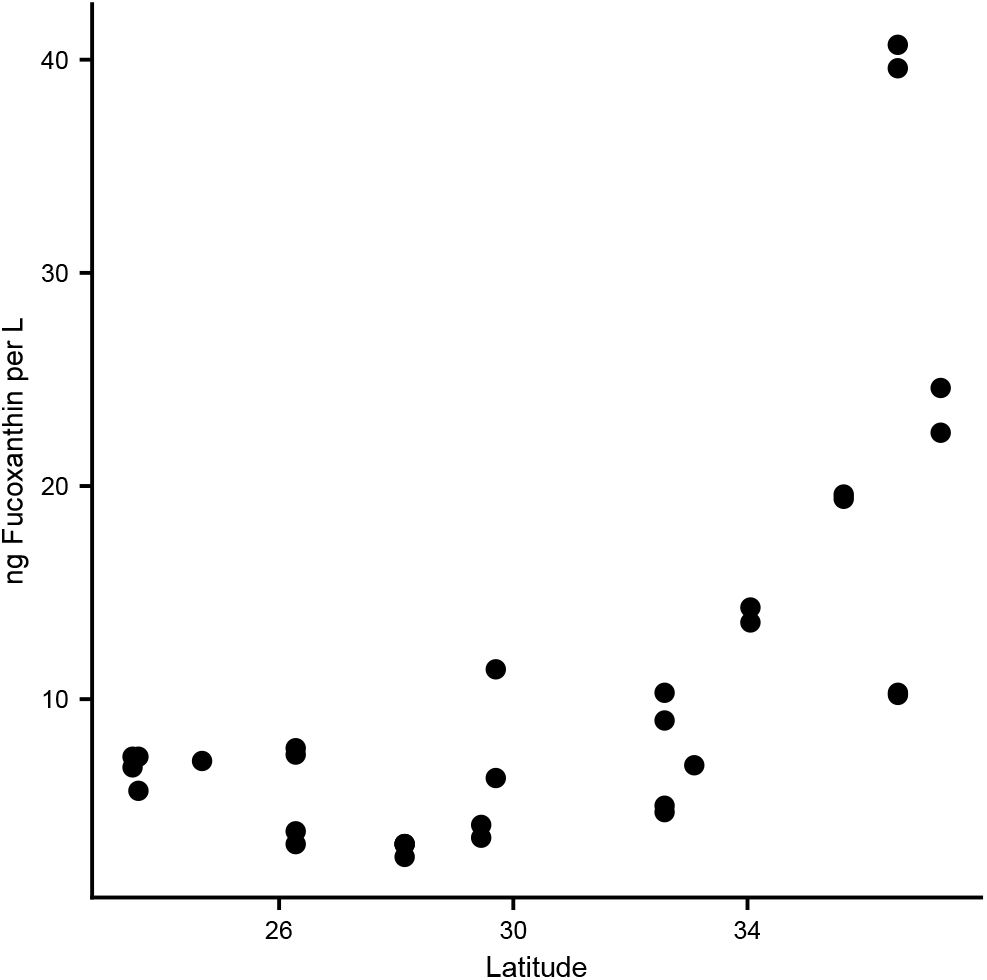
Concentration of the pigment fucoxanthin over latitude in transect samples.

**FIG S7.**
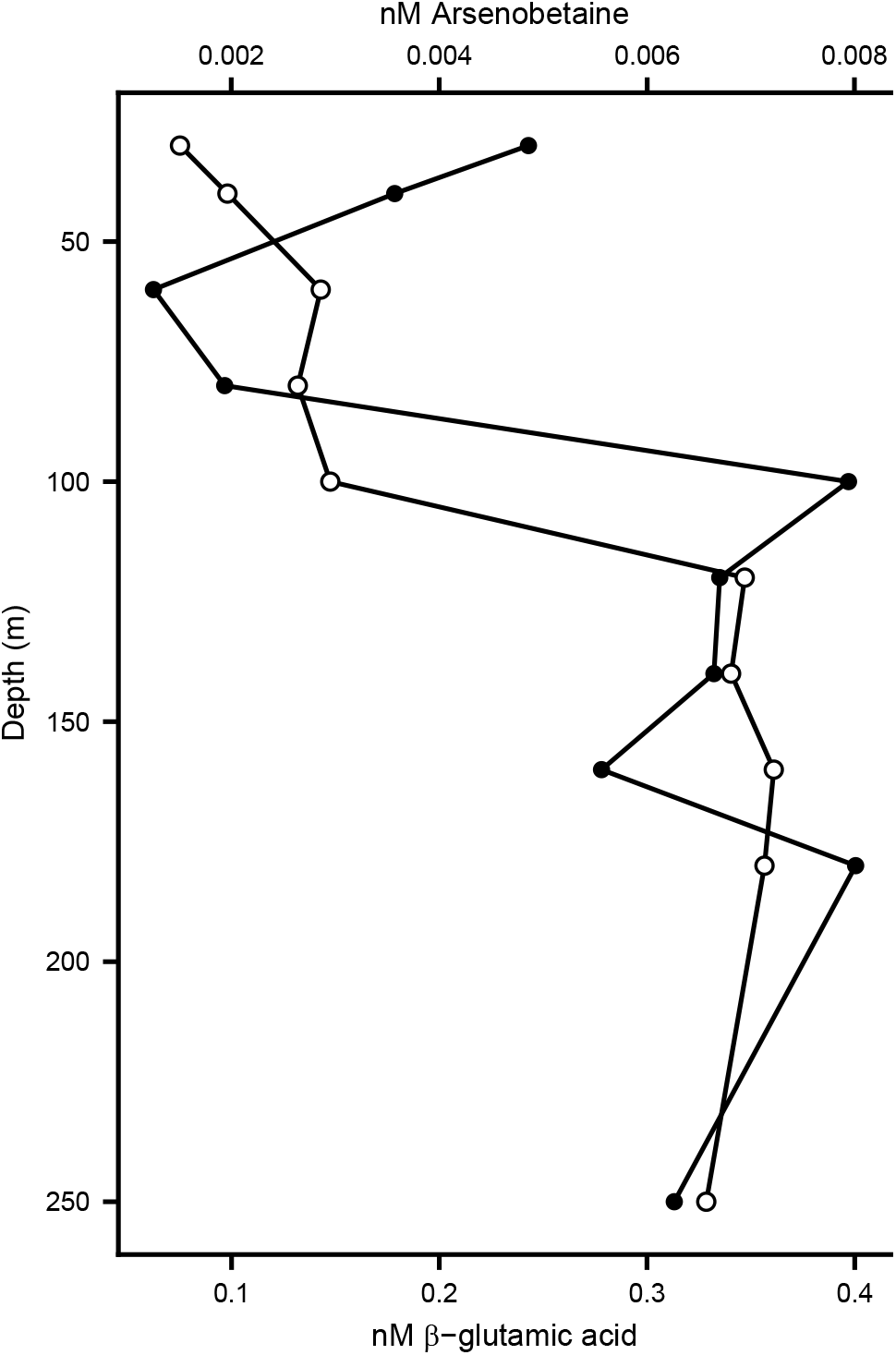
*β*-glutamic acid (closed circles) and arsenobetaine (open circles) depth profiles from the NPTZ. Note the different scales.

**FIG S8.**
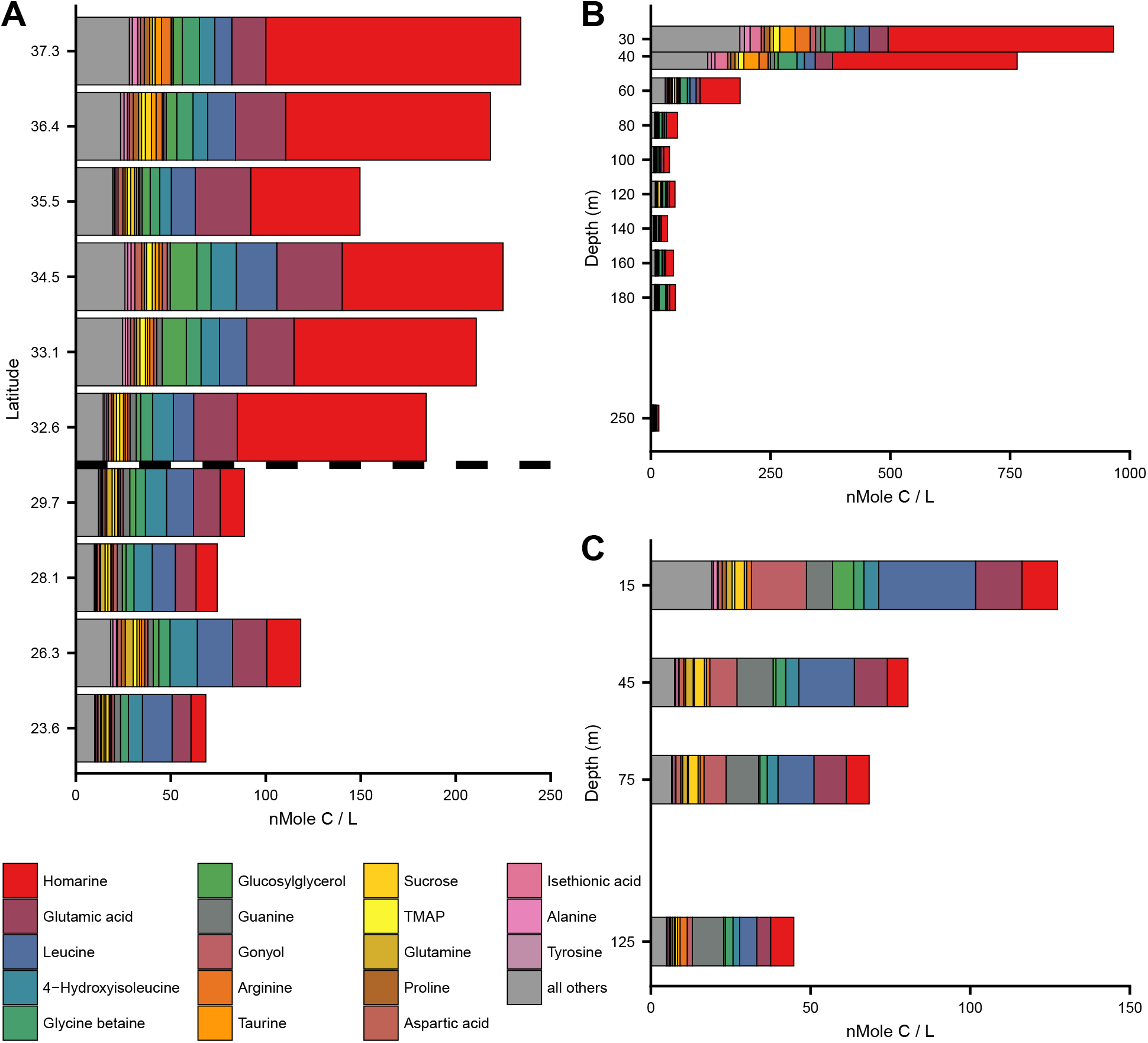
Most abundant 18 metabolites in environmental samples, presented as a nmole carbon per L. Meridional transet (A), with transition between NPSG to NPTZ shown as dotted line. Depth profile from NPTZ(B), and NPSG (C). Locations of samples are shown in Figure 1. These same data are presented as carbon mole fraction in Figures 4; full results in Table S9.

**FIG S9.**
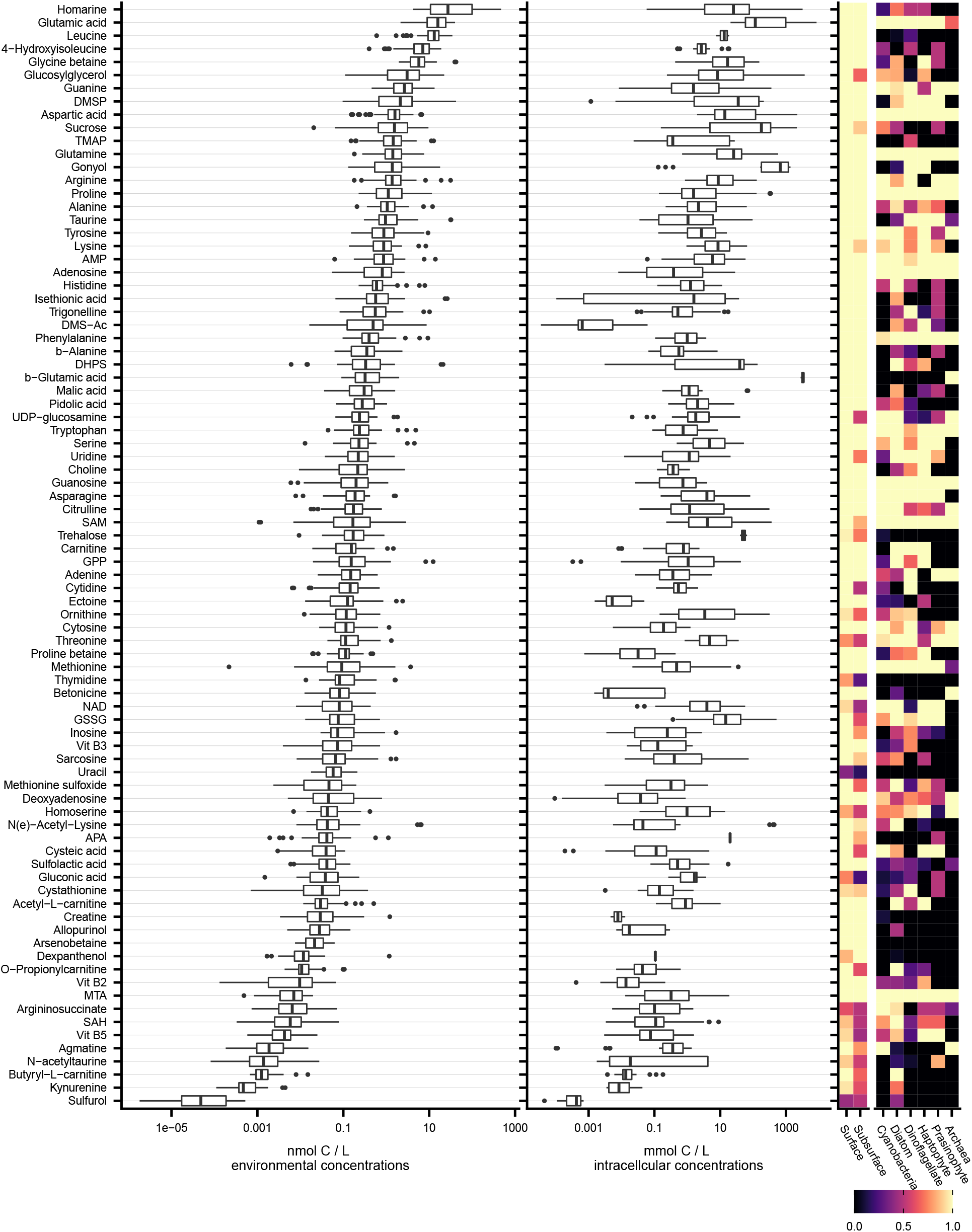
All identified compounds, with average and standard deviation (box and whiskers, respectively) of concentrations in surface seawater particles or in culture samples (excluding instances where we did not observe the compound). Right tiled panels shows the fraction of samples (environmental samples on the left, culture samples on the right) in which we observed these compounds. Note that x axis is on a log scale. Full results are found in Supplemental Table S10.

**FIG S10.**
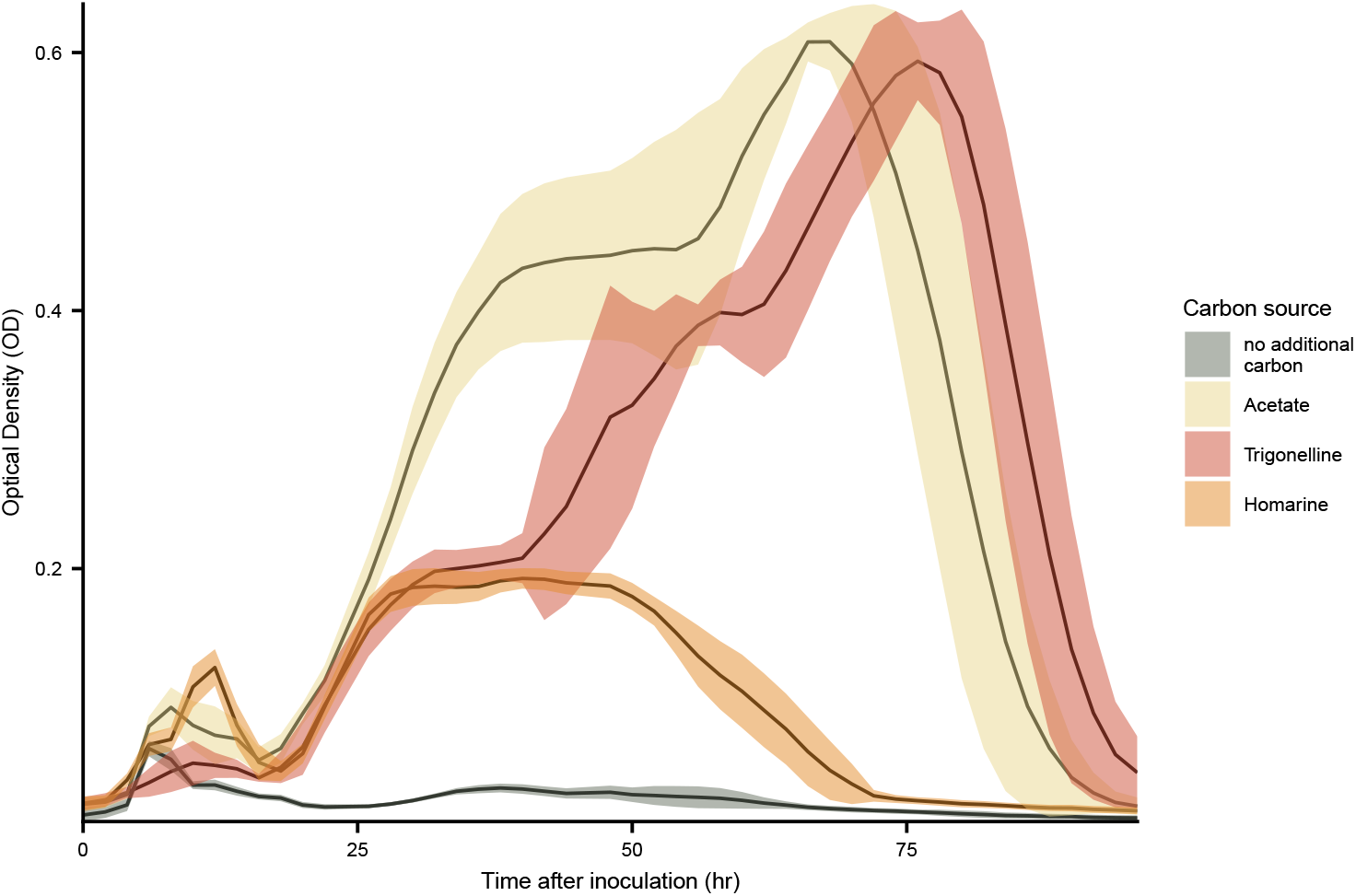
Growth curves of *Ruegeriapomeroyi* DSS-3 with three different carbon sources: acetate (positive control), homarine, and trigonelline, and no additional carbon (negative control). Note that the initial growth of the negative control is due to carryover of carbon from the inoculum, which had acetate as the carbon source. The same total carbon was added to each treatment.

**FIG S11.**
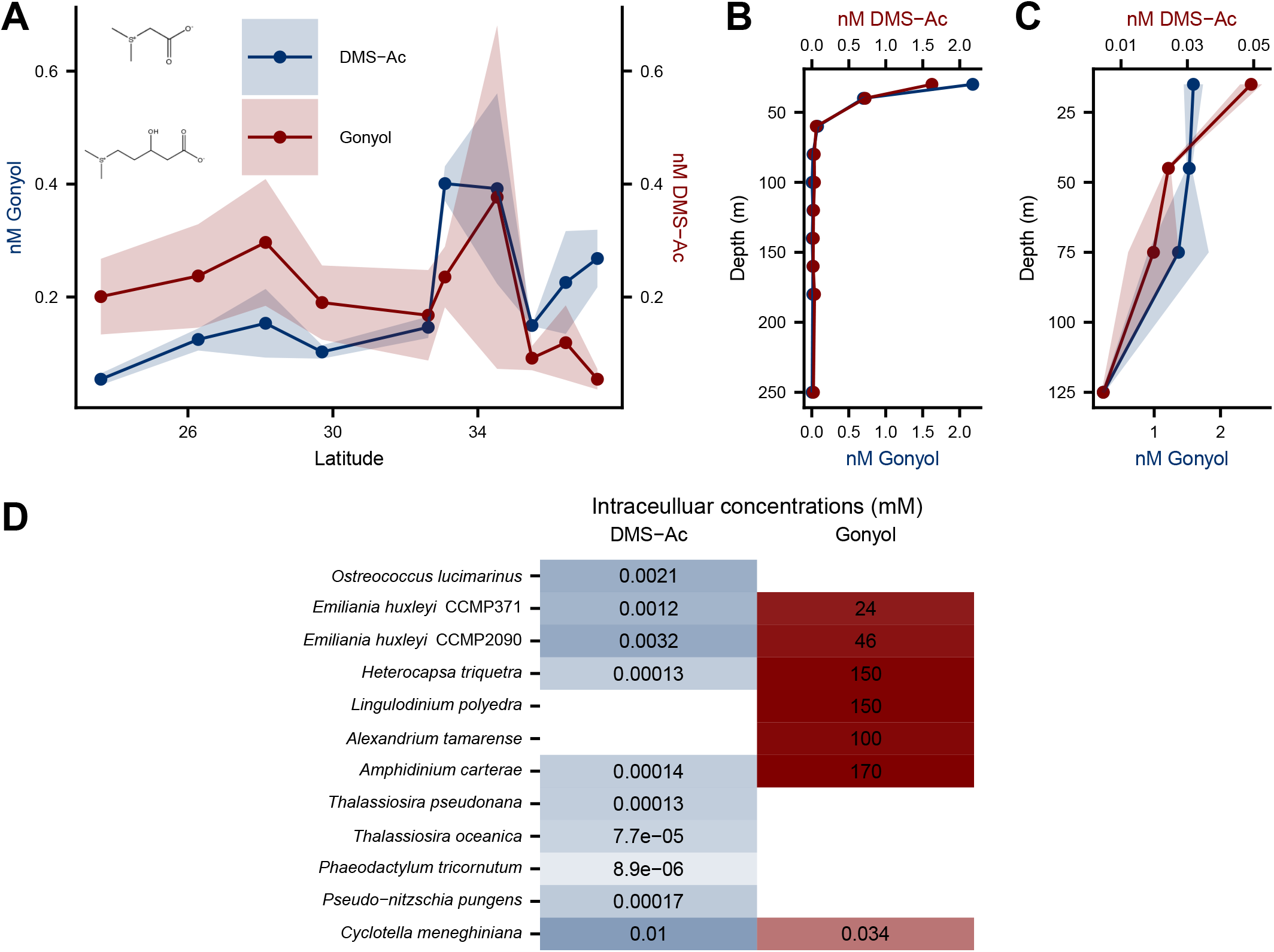
Gonyoland DMS-Ac spatial patterns in meridional transect (A), NPTZ depth profile (B), NPSG depth profile (C). Intracellular concentrations of gonyol and DMS-Ac in applicable organisms (E). Note the different scales for the compound concentrations (panel C) and depths (between panels B and C).

**TABLE S1.** Summary of samples collected and analyzed in this study.

| Cruise ID | Depth (m) | *n* | Date | Latitude | Longitude | Vol (L) | Collection method | Environmental regime |
| --- | --- | --- | --- | --- | --- | --- | --- | --- |
| KM1513 | 15.00 | 3 | 2015-07-31 | 24.55 | -156.33 | 11.00 | niskin | NPSG |
| KM1513 | 45.00 | 3 | 2015-07-31 | 24.55 | -156.33 | 9.00 | niskin | NPSG |
| KM1513 | 75.00 | 3 | 2015-07-31 | 24.55 | -156.33 | 13.00 | niskin | NPSG |
| KM1513 | 125.00 | 3 | 2015-07-31 | 24.55 | -156.33 | 12.00 | niskin | NPSG |
| KOK1606 | 15.00 | 2 | 2016-04-21 | 23.60 | -157.96 | 10.00 | niskin | NPSG |
| KOK1606 | 15.00 | 3 | 2016-05-01 | 26.28 | -158.00 | 10.00 | niskin | NPSG |
| KOK1606 | 15.00 | 5 | 2016-04-22 | 28.14 | -158.00 | 11.00 | niskin | NPSG |
| KOK1606 | 15.00 | 2 | 2016-04-23 | 29.45 | -158.04 | 15.00 | underway | NPSG |
| KOK1606 | 15.00 | 2 | 2016-05-01 | 29.70 | -158.00 | 10.00 | niskin | NPSG |
| KOK1606 | 15.00 | 2 | 2016-04-24 | 30.40 | -157.99 | 15.00 | underway | NPSG |
| KOK1606 | 15.00 | 5 | 2016-04-24 | 32.63 | -158.00 | 11.00 | niskin | NPTZ |
| KOK1606 | 15.00 | 2 | 2016-04-30 | 33.09 | -158.00 | 10.00 | niskin | NPTZ |
| KOK1606 | 15.00 | 2 | 2016-04-25 | 34.53 | -158.00 | 15.00 | underway | NPTZ |
| KOK1606 | 15.00 | 2 | 2016-04-26 | 35.49 | -158.01 | 13.00 | underway | NPTZ |
| KOK1606 | 15.00 | 2 | 2016-04-29 | 36.22 | -158.00 | 10.00 | underway | NPTZ |
| KOK1606 | 15.00 | 2 | 2016-04-29 | 36.30 | -157.99 | 10.00 | underway | NPTZ |
| KOK1606 | 15.00 | 2 | 2016-04-29 | 36.37 | -157.97 | 10.00 | underway | NPTZ |
| KOK1606 | 15.00 | 2 | 2016-04-29 | 36.46 | -157.96 | 10.00 | underway | NPTZ |
| KOK1606 | 15.00 | 2 | 2016-04-29 | 36.57 | -158.00 | 10.00 | niskin | NPTZ |
| KOK1606 | 15.00 | 4 | 2016-04-27 | 36.57 | -158.00 | 11.00 | niskin | NPTZ |
| KOK1606 | 15.00 | 2 | 2016-04-26 | 37.30 | -158.00 | 10.00 | niskin | NPTZ |
| MGL1704 | 30.00 | 1 | 2017-06-03 | 41.42 | -158.00 | 4.00 | niskin | NPTZ |
| MGL1704 | 40.00 | 1 | 2017-06-03 | 41.42 | -158.00 | 4.00 | niskin | NPTZ |
| MGL1704 | 60.00 | 1 | 2017-06-03 | 41.42 | -158.00 | 4.00 | niskin | NPTZ |
| MGL1704 | 80.00 | 1 | 2017-06-03 | 41.42 | -158.00 | 4.00 | niskin | NPTZ |
| MGL1704 | 100.00 | 1 | 2017-06-03 | 41.42 | -158.00 | 4.00 | niskin | NPTZ |
| MGL1704 | 120.00 | 1 | 2017-06-03 | 41.42 | -158.00 | 4.00 | niskin | NPTZ |
| MGL1704 | 140.00 | 1 | 2017-06-03 | 41.42 | -158.00 | 4.00 | niskin | NPTZ |
| MGL1704 | 160.00 | 1 | 2017-06-03 | 41.42 | -158.00 | 4.00 | niskin | NPTZ |
| MGL1704 | 180.00 | 1 | 2017-06-03 | 41.42 | -158.00 | 4.00 | niskin | NPTZ |
| MGL1704 | 250.00 | 1 | 2017-06-03 | 41.42 | -158.00 | 4.00 | niskin | NPTZ |

**TABLE S2.** Summary of physical and chemical parameters on April 2016 cruise. Reprinted with permission from (27).

| Region | Latitude (°N) | SST (°C) | SSS | N+N (μM) | PO<sub>4</sub> (μM) | Chl (mg m<sup>-3</sup>) |
| --- | --- | --- | --- | --- | --- | --- |
| NPSG | 23.54–29.7 | 19.8–24 | 35.1–35.3 | dl-0.002 | 0.02–0.05 | 0.01–0.11 |
| NPTZ | 32.63–37.3 | 11.4–17.1 | 34.1–34.7 | 0.06–5.87 | 0.07–0.51 | 0.16–0.37 |

**TABLE S3.** Summary of cultured organisms analyzed in this study. More information (including culturing conditions for all except the Archaea) can be found in (5). Cyanobacteria and archaea were obtained from individual lab culture collections and eukaryotic phytoplankton were obtained from the NCMA culture collection. More detailed information are in Table S7. Short ID is how the organism is labeled throughout the figures.

| Broad taxon | Species | Strain | *n* | Short ID |
| --- | --- | --- | --- | --- |
| Archaea | *Nitrosopumilus maritimus* | SCM1 | 3 | Nmar |
| Cyanobacteria | *Crocosphaera watsonii* | 8501 | 3 | 8501 |
| Cyanobacteria | *Prochlorococcus marinus* | 1314 | 3 | 1314P |
| Cyanobacteria | *Prochlorococcus marinus* | AS9601 | 3 | As9601 |
| Cyanobacteria | *Prochlorococcus marinus* | MED4 | 3 | MED4 |
| Cyanobacteria | *Prochlorococcus marinus* | NATL2A | 3 | Nat |
| Cyanobacteria | *Synechococcus sp.* | 7803 | 2 | 7803 |
| Cyanobacteria | *Synechococcus sp.* | 8102 | 2 | 8102 |
| Diatom | *Cyclotella meneghiniana* | 338 | 3 | Cy |
| Diatom | *Navicula pelliculosa* | 543 | 3 | Np |
| Diatom | *Phaeodactylum tricornutum* | 2561 | 2 | Pt |
| Diatom | *Pseudo-nitzschia pungens* | Pc55x | 3 | Pc55x |
| Diatom | *Thalassiosira oceanica* | 1005 | 3 | To |
| Diatom | *Thalassiosira pseudonana* | 1335 | 3 | Tp |
| Dinoflagellate | *Alexandrium tamarense* | 1771 | 3 | 1771 |
| Dinoflagellate | *Amphidinium carterae* | 1314 | 3 | 1314 |
| Dinoflagellate | *Heterocapsa triquetra* | 449 | 3 | 449 |
| Dinoflagellate | *Lingulodinium polyedra* | 2021 | 3 | 2021 |
| Haptophyte | *Emiliana huxleyi* | 2090 | 3 | 2090 |
| Haptophyte | *Emiliana huxleyi* | 371 | 3 | 371 |
| Prasinophyte | *Micromonas pusilla* | 1545 | 3 | 1545 |
| Prasinophyte | *Ostreococcus lucimarinus* | 3430 | 3 | 3430 |

